# Informing plasmid compatibility with bacterial hosts using protein-protein interaction data

**DOI:** 10.1101/2022.07.12.499728

**Authors:** Tim Downing, Min Jie Lee, Conor Archbold, Adam McDonnell, Alexander Rahm

## Abstract

The compatibility of plasmids with new host cells is significant given their role in spreading antimicrobial resistance (AMR)^1^ and virulence factor genes. Evaluating this using *in vitro* screening is laborious and can be informed by computational analyses of plasmid-host compatibility through rates of protein-protein interactions (PPIs) between plasmid and host cell proteins. We identified large excesses of such PPIs in eight important plasmids, including pOXA-48, using most known bacteria (n=4,363). 23 species had high rates of interactions with four *bla*OXA-48-positive plasmids. We also identified 48 species with high interaction rates with plasmids common in *Escherichia coli*. We found a strong association between one plasmid and the fimbrial adhesin operon *pil*, which could enhance host cell adhesion in aqueous environments. An excess rate of PPIs could be a sign of host-plasmid compatibility, which is important for AMR control given that plasmids like pOXA-48 move between species with ease.

**Highlights:**

- We tested for protein interactions between key plasmids and 4,363 bacterial hosts
- 23 host species had high rates of protein interactions with four *bla*OXA-48 plasmids
- *Pseudomonas* species showed high rates of interactions with these plasmids
- Host-plasmid protein-protein interaction rates may be associated with compatibility

## 1. Introduction

Plasmids can provide new genes to host cells, enhancing properties like antimicrobial resistance (AMR) and adhesion to host molecules [1,2]. Plasmids are short extrachromosomal DNA elements that are usually circular, such that conjugative plasmids can move between different host cells sharing the same ecological niche [3,4]. DNA transformation, extracellular vesicles and bacteriophages also permit plasmid gene transfer, albeit less often than conjugation [5-8]. Collectively, these horizontal gene transfer (HGT) processes meaning that the role of plasmids in enabling adaptation to new environments is a key concern in addressing the spread of infectious bacteria. Additionally, plasmids typically induce a small fitness cost in their host cells [9,10]. To overcome this, many plasmids possess genes to parasitise host cells via post-segregation killing and stable inheritance: this ensures their propagation in daughter cells [11] and facilitates vertical plasmid transmission. Many plasmids also endow their host cells with enhanced AMR [12], thereby enhancing their fitness and subsequently the plasmid’s survival. Consequently, clonal radiations of emerging pathogens are often associated with the HGT-mediated acquisition of plasmids encoding key AMR genes (like *bla*OXA-48) that enable survival in antibiotic-rich niches.

Plasmid-encoded AMR genes help drive the transmission of carbapenemase-producing *Enterobacteriaceae* (CPE) including *Escherichia coli* and *Klebsiella*. These genes are concerning because they can be transmitted to other bacteria. These two priority pathogens impair human health according to the World Health Organisation (WHO) [13] and were the most common CPEs in England in 2014-16 at 27% (*E. coli*) and 52% (*Klebsiella*) where the most common carbapenemase gene was *bla*OXA-48 [14]. More recent surveys of CPEs in England in 2020-22 confirmed this pattern, showing *E. coli* and *Klebsiella* were responsible for 26% and 33% of cases, respectively [15]. Such plasmid-driven AMR gain by extraintestinal pathogenic *E. coli* (ExPEC) has enhanced their capacity to infect epithelial cells beyond their normal gut niche [16-25].

Highly transmissible pOXA-48-like plasmids are a compelling example of challenges driven by mobile genetic elements [26]. The *bla*OXA-48 gene encodes an Ambler class D beta-lactamase first identified in *Klebsiella pneumoniae* from Turkey [27]. It is usually on a highly mobile 62 Kb conjugative plasmid named pOXA-48a [28] that has spread the *bla*OXA-48 gene globally, causing substantial outbreaks in many countries, including Ireland [29-32]. An estimated 87% of *bla*OXA-48-positive isolates are MDR, and about half of CPE isolated in Europe have this group II OXA carbapenemase gene [33], including *Serratia maracescens* (since 2021). This gene has been found regularly in community and hospital settings across diverse host species, most typically in the *Enterobacterales* order. Host-pOXA-48a co-evolution has occurred, for example through host cell wall permeability changes to alleviate an initial low level of *bla*OXA-48 carbapenemase activity [34].

Beyond pOXA-48, IncF plasmids’ roles in spreading AMR genes are well known [12,35-36], including non-conjugative plasmids pEK499 and pEK516 in MDR *E. coli* [37-40]. These plasmids were first identified in England in *E. coli* and have ∼75% similarity to each other [37]. Additionally, a distinct conjugative IncI1 *bla*CTX-M-3-*bla*TEM-1b-positive plasmid pEK204 was found in epidemiologically related hosts in Belfast [37-38]. Plasmid pEK204 encodes a type IV pilus (T4P) through its *pil* operon which encodes a thin pilus for epithelial cell adhesion or biofilm formation [41]. Plasmids encoding such fimbrial adhesins could allow bacterial cells to survive in new environments because adherence to human epithelial cells facilitates bacterial colonisation and the formation of biofilms [42]. These adhesins include type 1, P, F1C/S and AFA fimbriae/pili that are typically assembled by the chaperone-usher secretion pathway in gram-negative bacteria [43-47]. Certain pathogens retain multiple different fimbriae that can be expressed depending on the environmental conditions [48-49]. This indicates that possessing a repertoire of fimbriae could be adaptively beneficial. T4P are thin connections that can be elongated or retracted by ATPases [50] and are associated with biofilm formation [51], virulence [52] and bacteriophage infection [53]. The *pil* operon (particulary *pilS*) may allow differential epithelial cell or abiotic surface adhesion with a thin flexible pilus [41] that can elevate conjugation rates in liquid environments [54]. It can also permit biofilm formation such that a T4P-negative cell can adhere to other T4P-expressing cells bound with a host cell [41]. Although the *pil* operon is somewhat rare, its widespread nature across bacteria implies a role in human infection [55].

These and other findings imply that a plasmid’s capacity to interact successfully with a host cell is important and stems more from its genetic profile more so than the host genome [4]. This suggests host fitness increases following plasmid gain depend not only on the environment, but also on plasmid-host protein-protein interactions (PPIs) arising from their collective genetic profiles. These PPIs must be beneficial for plasmids to persist because plasmid-encoded genes must be replicated and expressed in the host cell. This presents the hypothesis that higher rates of plasmid-host PPIs are a signature of higher compatibility between the bacterial cell and the plasmid.

Limited work has examined plasmid-host PPIs across all bacteria. Previously, PPI network analysis has identified AMR genes in *E. coli* [55,56], but has not been applied at the level of whole plasmids. Here, we examined PPIs between plasmid and bacterial hosts to explore the potential compatibility between the host cell and plasmid. We focused on four pOXA-48-related isoforms and four *E. coli* plasmids (pEK204, pEK499, pEK516, pEC958). We assessed these features initially in 4,363 samples representative of most bacteria. We found that it was possible that elevated sample-plasmid PPI levels could be symptomatic of host-plasmid compatibility, and that this observation may stem from operons associated with environments where those bacterial samples have been found.

## 2. Methods

### 2.1 Protein-protein interaction data extraction for 24 *Enterobacterales* and all bacteria

We retrieved PPI information for two key datasets from STRING database v12 [57] using R packages Rentrez v1.2.2 [58], STRINGdb v2.0.1 [59], and StringR v1.4.0 [60] (Figure 1), in a working environment using Rv4.0.1 [61], RStudio v2022.2.3.492 [62], Bioconductor v3.11 [63]. The dataset first was for three genera from the *Enterobacterales* order containing: *Escherichia* and *Shigella* (n=11), *Klebsiella* (n=2) and *Serratia* (n=11) (Table S1). *Serratia* belongs to the *Yersiniaceae* family, whereas the others are all *Enterobacteriaceae*. Three other potential samples in StringDB belonging to these genera were excluded because they lacked any pOXA-48-related plasmid-sample PPIs and so had no information. The second dataset was composed of 4,363 StringDB bacterial samples (including the 24 *Enterobacterales* from above) with valid PPI data as outlined previously [79] (Figure 1). Using a score threshold of 400 as per previous work [79], the number of proteins and their PPIs were recorded. These StringDB scores come from a range of sources, including: text mining, pathway databases, co-expression research, small-scale and high-throughput experiments, and computational predictions (chromosomal proximity, phylogenetic context and gene fusion) [57]. The data was processed, collated and visualised using Dplyr v1.0.8 [64], Forcats v0.5.1 [65], Ggplot2 v3.3.5 [66], Ggrepel v0.9.1 [67], ReadR v2.1.2 [68], Readxl v1.4.0 [69], Tibble v3.1.6 [70], TidyR v1.2.0 [71], Tidyverse v1.3.0 [72] and VennDiagram v1.7.3 [73].

**Fig 1.**
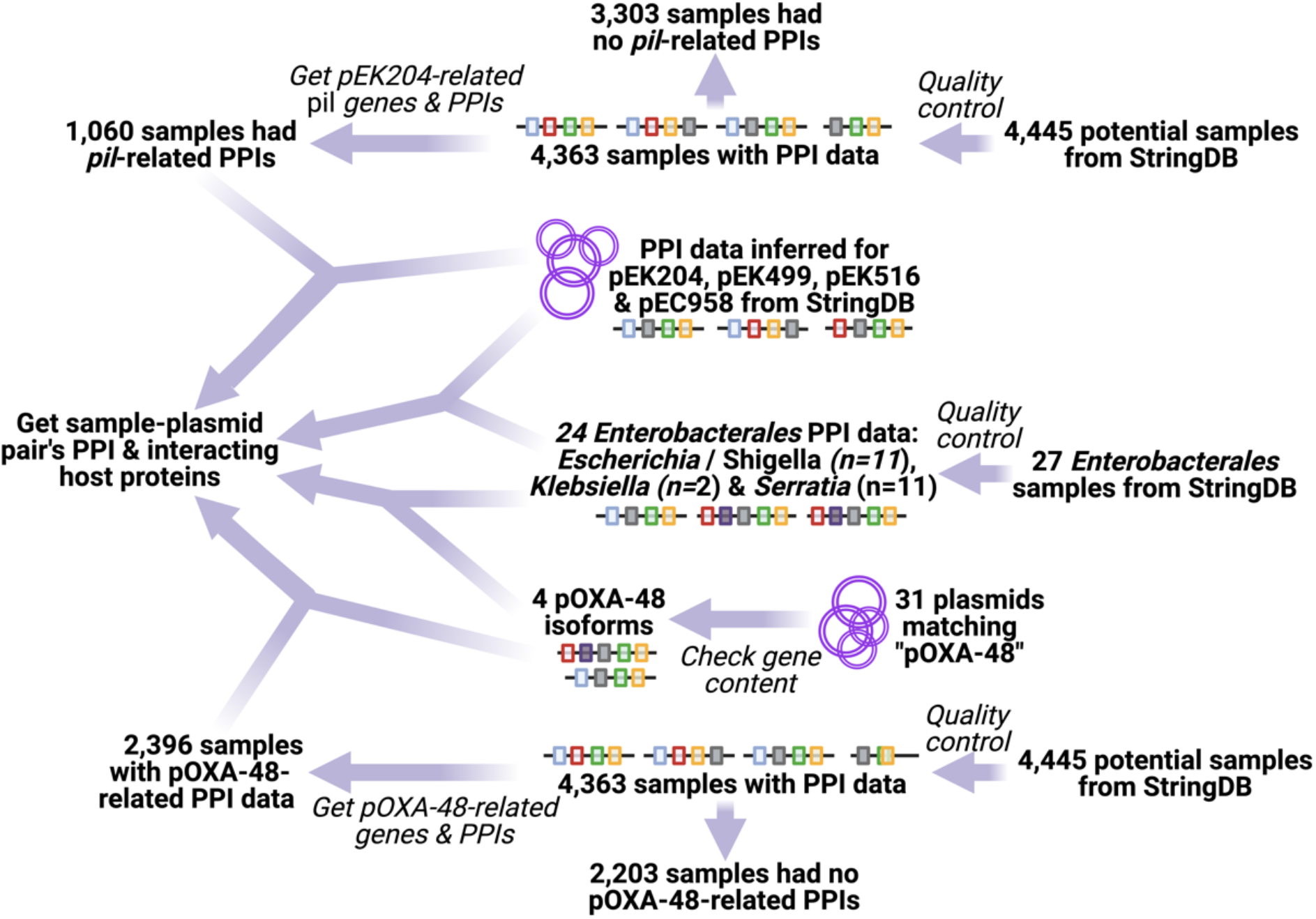
The data sources and methods summary, read from right to left. We used PPI data from StringDB related to two datasets: one with 24 *Enterobacterales* (middle) and another with 4,363 diverse bacterial samples (top and bottom). Top: We compared the interacting host proteins and associated PPIs for four plasmids pEK204, pEK499, pEK516 and pEC958 compared to the 24 *Enterobacterales* samples’ PPI data to assess their PPIs. These plasmids were also compared to the 4,363 samples’ PPI data. Bottom: We examined 31 initial pOXA-48-related plasmids that resolved as four main isoforms whose interacting host proteins and associated PPIs we compared to the 24 *Enterobacterales’* PPI and the 4,363 samples in the same way. Created with BioRender.com.

### 2.2 Plasmid data retrieval and quality control

The IncL pOXA-48a plasmid has been strongly associated with *E. coli* and *K. pneumoniae* [28]. To explore this plasmid type further, we downloaded accession data for all (n=31) NCBI NucCore entries containing “pOXA-48”. Plasmid pKP112 possessed a *bla*OXA-48 gene and therefore was considered a valid pOXA-48 plasmid in this case. These came from the three *Enterobacterales* species: *E. coli* (n=7), *K. pneumoniae* (n=22), and *Serratia marcescens* (n=2). Among these 31 sequences, there were four genetically distinct pOXA-48 isoforms, all isolated from *K. pneumoniae* (strains named Kp11978, KP112, Kpn-30715/15 and K8) (Table 1). Among these four isoforms, pOXA-48_3071 had a *bla*OXA-48 gene that was not properly annotated in the GenBank entry, so it was considered as a pOXA-48-positive plasmid here. We also obtained the unique proteins of four important plasmids linked to antibiotic resistance in *E. coli*: pEK204 (IncI1), pEK499 (IncFII-FIA), pEK516 (IncF) and pEC958 (IncF) (Table S2). Plasmids pEC958 and pEK499 have high similarity (∼92%) [74].

**Table 1.** The plasmids examined came from two groups. The four pOXA-48-related plasmids were originally identified in *K. pneumoniae* (Strain of origin), and the four linked to *Enterobacterales* were initially isolated in *E. coli*.

| Plasmid group | Plasmid name | Strain of origin | Accession number | Length (bp) | Genes | CDSs |
| --- | --- | --- | --- | --- | --- | --- |
| pOXA-48 | pOXA-48 | Kp11978 | JN626286.1 | 61,881 | 40 | 74 |
|  | pKP112 | KP112 | LN864819.1 | 84,252 | 62 | 102 |
|  | pOXA-48_30715 | Kpn-30715/15 | NZ_KX523901 | 65,488 | 84 | 84 |
|  | pOXA-48_k8 | K8 | MT441554.1 | 65,499 | 87 | 87 |
| *Entero-bacteriales* | pEK204 | Strain C | EU935740 | 93,732 | 89 | 95 |
|  | pEK499 | Strain A | EU935739 | 117,536 | 103 | 141 |
|  | pEK516 | Strain D | EU935738 | 64,471 | 65 | 79 |
|  | pEC958 | EC958 | NZ_HG941719 | 135,602 | 153 | 153 |

To assess plasmid protein similarity, we measured the amino acid sequence similarity of the translated genes of the eight plasmids in a pairwise manner using R package Biostrings v2.60.2 [75]. Among the proteins shared by the pOXA-48-related plasmid pairs, 94% (132 out of 138) had a match (*bitscore <* −*100*). Among the proteins shared by pEK499, pEK516 and pEC958, 94% (130 out of 140) had a match, but pEK204’s comparisons with pEK499, pEK516 and pEC958 showed less similarity (18%). Comparisons between each group (pOXA-48-related vs pEK204, pEK499, pEK516 and pEC958) showed only 22% of protein-level similarity for shared proteins.

The PPI information for these eight plasmids was compared with StringDB PPI data for the 24 *Enterobacterales* and 4,363 bacterial samples above using the above packages in R v4.0.1 and Genbankr v1.20.0 [76]. We computed the number of host proteins interacting with each plasmid protein, the number of host-plasmid PPIs, and the Jaccard index of the fraction of interacting host proteins compared to the combined total of sample and plasmid proteins. The number of genes per plasmid was recorded and analysed using R package UpSetR [77]. All p values reported in this study were corrected for multiple testing using the Benjamini-Hochberg method in R.

### 2.3 Plasmid comparison and estimation of an excess of protein-protein interactions

2,021 samples (out of 4,363) had PPIs with at least one of these four pOXA-48 isoforms, of which 650 had PPIs with all four isoforms. Samples without PPIs were not examined because these provided no information. There were 6,181 sample-plasmid pairs in total with ≥10 interacting proteins and ≥10 PPIs, coming from: pOXA-48 (1,820 pairs), pKP112 (1,868), pOXA-48_30715 (673) and pOXA-48_k8 (1,820). For the four *E. coli* plasmids (pEK204, pEK499, pEK516 and pEC958), all 24 *Enterobacterales* and 1,604 other samples had ≥1 PPI with at least one of these plasmids. There were 3,828 sample-plasmid combinations with ≥10 interacting host proteins and ≥10 PPIs across these samples, coming from: pEK204 (856), pEK499 (1,324), pEK516 (852) and pEC958 (796).

To quantify instances of high host-plasmid PPI rates for each plasmid set above, we examined on the association between the numbers of interacting host proteins per sample and the number of sample-plasmid PPIs using a linear model predicting the number of PPIs per sample-plasmid pair. We based the prediction on the numbers of: interacting host proteins, plasmid proteins, proteins per sample, and associated pairwise interactions for each plasmid set (Table S3). This showed that the only informative variables were the number of interacting proteins and the number of PPIs per sample and not the other variables. Thus, we used these parameters in a linear model to get the residual numbers of PPIs per sample-plasmid pair, reflecting instances where the observed number of PPIs was far in excess of the expected number. These residuals from the resid R function were normalised such that the average excess of PPIs was zero. We did not explore deficits in the observed PPI numbers.

### 2.4 Interactions between fimbrial adhesin operons

We focused further on pEK204’s *pil* operon, which is associated with adhesion and has 14 genes, *pilI-V* [37]. We used pEK499 as a baseline because it had the most samples with PPIs (1,324 out of 4,363) among the other three plasmids. We found 1,060 samples had detected PPIs with both pEK204 and pEK499. We examined them using a normalised log2-scaled ratio of the Jaccard indices of pEK204 versus pEK499. The pairwise Pearson correlation between 14 *pil* genes was determined using ppcor v1.1 [78] based on presence-absence data for 4,377 samples from across bacteria [79]. Genes with multiple synonyms were collapsed into a single representative. These 14 *pil* genes were also compared to 14 *fim* genes and four *pap* genes in the same manner. We retrieved the *pil, fim* and *pap* gene synonyms and associated Pil, Fim and Pap PPIs in the gene name and PPI data for the 4,363 samples.

## 3. Results

To test for excess PPI rates with different hosts and plasmids, we compared the PPIs of four pOXA-48 plasmid isoforms and four key *E. coli* plasmids with two sample collections whose PPI data was retrieved from StringDB. The first sample collection consisted of 24 *Escherichia, Klebsiella, Serratia* and *Shigella* samples to examine genus-level plasmid links, and the second collection had 4,363 samples from across bacteria to inform on wider patterns.

### 3.1 No genus-level association of pOXA-48 within *Enterobacterales*

We examined the PPI levels of four known pOXA-48 isoforms (pOXA-48, pKP112, pOXA-48_30715, pOXA-48_k8, Table 1) with 24 *Enterobacterales* from *Escherichia* and *Shigella* (n=11), *Klebsiella* (n=2) and *Serratia* (n=11) (Table S1). There was a wide range of PPI levels but no genus-level association for any of these plasmids, consistent with previous work [80]. To resolve this finding, we examined the genetic composition of these plasmids. These pOXA-48 isoforms shared just four genes including *ssb, pemI/K* and *bla*OXA-48 (Figure 2A) (*bla*OXA-48 was present but not annotated on pOXA-48_3071). PemI is the antitoxin of a type II toxin-antitoxin (TA) system for post-segregational killing that can neutralise its endoribonuclease toxin, PemK [81]. These differences suggested that the plasmids’ effects on host cell PPI networks would vary.

**Fig 2.**
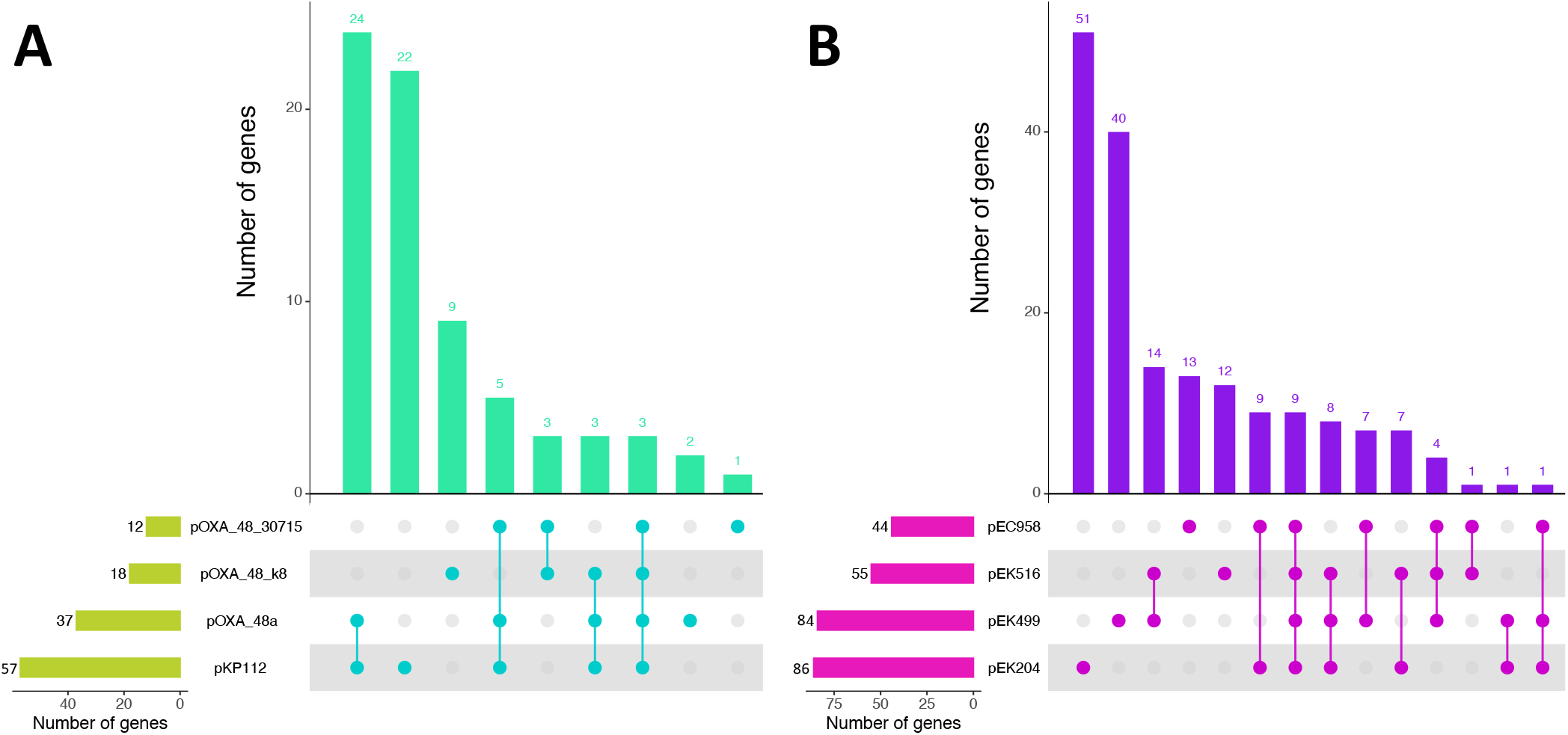
(A) The overlap of unique annotated genes between four pOXA-48-related plasmids (pOXA-48, pKP112, pOXA-48_30715, pOXA-48_k8). Top: The intersection sizes (y-axis) and the genes per set (green) showed that only four genes were shared by all four plasmids: *ssb, pemI/K* and *bla*OXA-48 (*bla*OXA-48 was not annotated properly on pOXA-48_3071 and so the intersection of all four plasmids shows three genes but contains *bla*OXA-48 too). Bottom: The numbers of genes per sample (left, lime-green, set size) with the corresponding sets (right: cyan circles). The most divergent isoform was pKP112, which had 22 unique genes. (B) The overlap of genes between the four common *E. coli* plasmids (pEK204, pEK499, pEK516, pEC958). Top: The intersection sizes (y-axis) and the genes per set (purple) showed that nine genes were shared by all four plasmids: *ssb, psiB* and *traA*/*E*/*L*/*K*/*R*/*V*/*Y*. Bottom: The numbers of genes per sample (left, pink, set size) with the corresponding sets (right: purple circles). The most divergent isoform was pEK204 (51 unique genes).

### 3.2 Incomplete associations of pEK204, pEK499, pEK516 and pEC958 with *Escherichia* species

Next, we assessed the level of PPIs between these 24 *Enterobacterales* and four major plasmids (pEK204, pEK499, pEK516, pEC958, Table 1). These plasmids possess numerous AMR genes and are prevalent, particularly in certain *E. coli* phylogroups. As above, the numbers of interacting host proteins per sample and PPIs had no clear association (Table S1). Again, there were elevated rates of interacting proteins and PPIs for nine of ten *Escherichia* samples, but *Klebsiella* and *Serratia* were more heterogeneous (Table S1). Consequently, the plasmids’ proteomes appeared to be the major factors, like pOXA-48. The gene content of pEK204, pEK499, pEK516 and pEC958 showed nine shared genes: *ssb, psiB* and *traA*/*E*/*L*/*K*/*R*/*V*/*Y* (Figure 2B). The *tra* region is associated with conjugation, *psiB* encodes a plasmid SOS inhibition protein that stops induction of the SOS pathway by binding RecA [82], and *ssb* encodes a ssDNA-binding protein involved in DNA replication, recombination and repair [83].

### 3.3 Varied patterns of pOXA-48 interactions across bacteria

To shed light on these complex patterns, we extended our analysis of the four pOXA-48 plasmids to all bacterial samples with valid PPI data from StringDB (n=4,363), focusing on 6,181 sample-plasmid combinations with ≥10 interacting host proteins and ≥10 PPIs. Plasmid pOXA-48_30715 had the most interacting host proteins (53±23, median±SD) and PPIs (53±26), followed by pKP112 (interacting proteins, 39±45; PPIs, 43±65), pOXA-48 (interacting proteins, 35±45; PPIs, 37±58), and then pOXA-48_k8 (interacting proteins, 30±33; PPIs, 31±37) (Figure S1). Notably, the interacting host proteins per plasmid had the largest association with the number of PPIs per pair (0.957 ≤ r ≤ 0.995, Figure S2).

Only 0.6% (40 out of 6,401) of the sample-plasmid pairs had high levels of association, measured as an excess of plasmid-linked PPIs ≥100, ≥100 interacting host proteins and ≥20% excess PPIs after using a model to correct for the number of interacting host proteins per sample-plasmid pair (Figure S3, Table S4). 22 of these 40 were linked to pKP112, 13 to pOXA-48, three to pOXA-48_k8 and two to pOXA-48_30715 (Figure 3). *P. aeruginosa* had the most interacting host proteins of these 40: it shared 19 proteins on pKP112 (Figure S4) that had PPIs with 450 other *P. aeruginosa* proteins (Figure S5) with 684 PPIs, an excess of 27%. *P. aeruginosa* also had >300 interacting host proteins, >340 PPIs and an excess of PPIs for pOXA-48. The sample with the 2^nd^ highest number of interacting host proteins was *Aromatoleum aromaticum* EbN1, which had 32 proteins on pOXA-48 (Figure S6) interacting with 252 *A. aromaticum* proteins with 400 PPIs, an excess of 34% (Figure S7). This sample also had 297 interacting proteins linked to pKP112 with 452 PPIs, an excess of 28%. *Enterobacter cloacae subsp. cloacae* ATCC 13047 had 125 other host proteins interacting with pKP112 (Figure S8) with 173 PPIs (an excess of 20%, Figure S9). Two *Legionella* and a *Citrobacter* species also had high associations with pKP112 and pOXA-48. *Klebsiella oxytoca* had 173 interacting host proteins with pKP112 that had 221 PPIs (an excess of 9%), but no such association with the other three plasmids.

**Fig 3.**
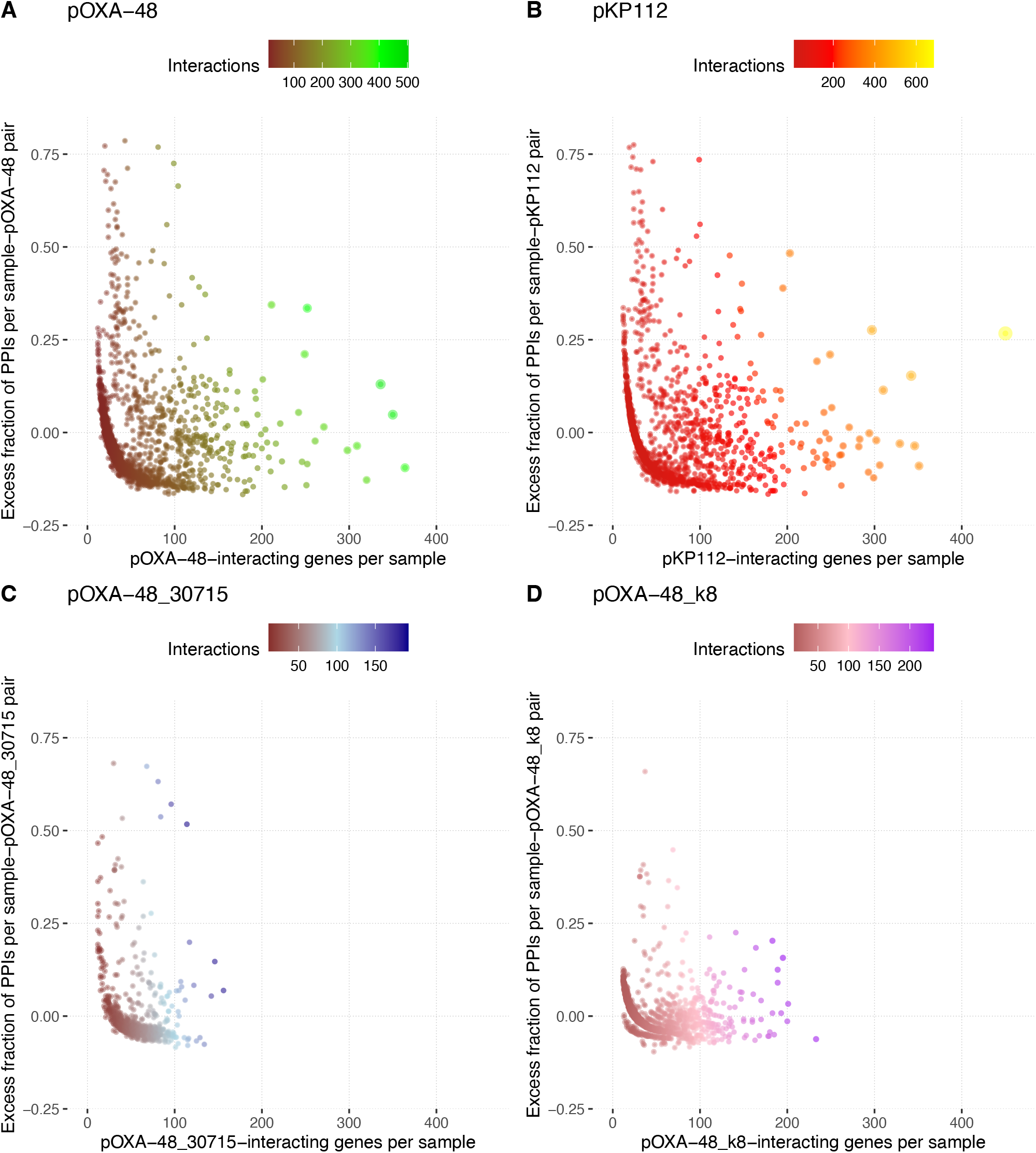
The numbers of interacting host proteins per sample (x-axis) versus the excess fraction of PPIs (y-axis) for each sample-plasmid pair for the four pOXA-48 isoforms (A) pOXA-48 (red-green), (B) pKP112 (red-yellow), (C) pOXA-48_30715 (brown-blue) and (D) pOXA-48_k8 (mauve-purple) using data from 6,181 samples’ plasmids with >10 interacting host proteins and >10 PPIs each. The number of PPIs per sample-plasmid pair is denoted by the point size and by the shade based on the scale: larger means fewer PPIs, smaller means more PPIs. Note this scale differs for each plasmid. In (B), *P. aeruginosa* had the most interacting proteins with pKP112 (yellow).

### 3.4 Varied protein-level interactions of bacteria with pEK204, pEK499, pEK516 and pEC958

Assessing pEK204, pEK499, pEK516 and pEC958 in the same manner with the larger collection of 4,363 samples, we identified 1,604 samples with ≥1 PPI with at least one of these plasmids. We focused further on 3,828 sample-plasmid pairs with ≥10 interacting host proteins and ≥10 PPIs in 1,410 samples, of which 48.4% (683 out of 1,410) had PPIs with all four plasmids. The number of interacting host proteins per sample was comparable across all four plasmids, and highest for pEK204 (median of 64 vs 54 for the others, Figure S10). The distributions of numbers of PPIs per sample were similar, but with a much higher variance for pEK204 relative to the other three (13.8 vs 8.2), highlighting that samples had elevated PPI rates (Figure S11).

Here, 1.5% (56 out of 3,828) of sample-plasmid pairs had strong associations, measured as an excess of plasmid-linked PPIs ≥300, ≥100 interacting host proteins and an excess of PPIs of ≥20% (Table S5). 48 were associated with pEK204, five with pEC958, two with pEK499, and one with pEK516 (Figure 4). In particular, the number of pEK204 PPIs per pair varied much more than the other plasmids (Figure S12). Nine of the total of 56 sample-plasmid pairs were from the *Pseudomonas* genus, a pathogen existing in diverse environments [84]. *P. fluorescens* F113 (aka *P. ogarae* F113) had the most interacting host proteins with pEK204 (500) linked to 1,215 PPIs, an excess PPI rate of 22%. As with pKP112 and pOXA-48, *P. aeruginosa* had 24 pEK204-encoded proteins (including 12 from *pil*) (Figure S13) interacting with 596 other *P. aeruginosa* proteins (Figure S14) with 1,187 PPIs (an excess of 4%). Like pKP112 and pOXA-48 too, *A. aromaticum* EbN1 had 44 proteins on pEK204 including those from the *pil* operon and *tra* region (Figure S15) interacting with 582 other *A. aromaticum* EbN1 proteins (Figure S16) with 1,191 PPIs (an excess of 6%).

**Fig 4.**
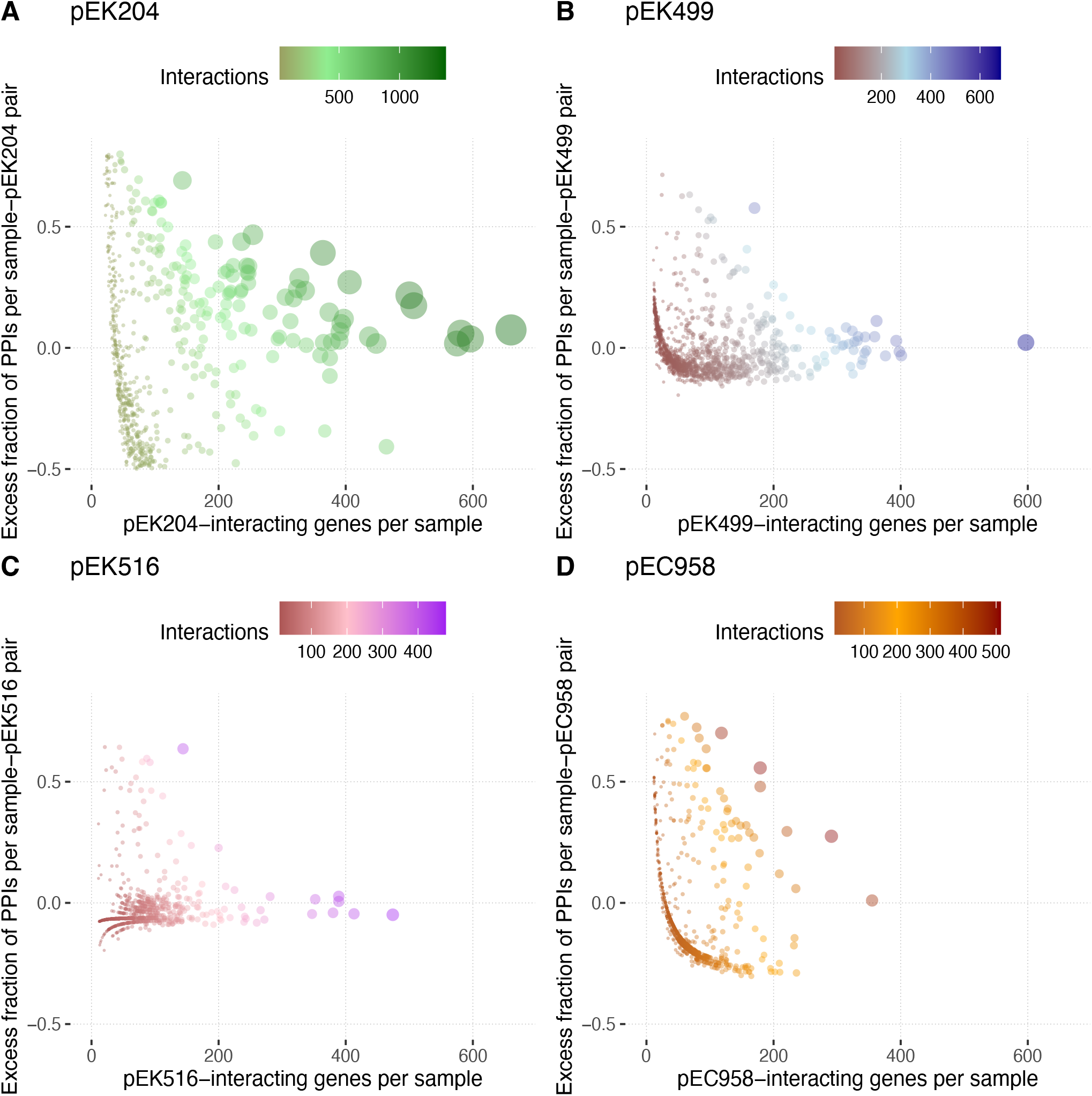
The numbers of interacting host proteins per sample (x-axis) versus the excess fraction of PPIs (y-axis) for each sample-plasmid pair for (A) pEK204 (brown-green), (B) pEK499 (brown-blue), (C) pEK516 (red-mauve) and (D) pEC958 (orange-brown) using data from 3,828 samples’ plasmids with 10+ interacting proteins and 10+ PPIs each. The number of PPIs per sample-plasmid pair is denoted by the point size and by the shade based on the scale: larger means fewer PPIs, smaller means more PPIs. Note this scale differs for each plasmid. In (A) for pEK204, the five samples with the most interacting proteins per sample were *P. fluorescens* F113 (aka *P. ogarae* F113), *P. aeruginosa, Aromatoleum aromaticum* EbN1, *Rubrivivax gelatinosus* IL144 and *Cellvibrio* sp. BR were the six rightmost samples (all green). *E. coli* K12 MG1655 had the most interacting proteins with pEK499 (B, blue) and pEK516 (C, mauve). In (D), *Ralstonia eutropha* H16 had the most interacting proteins with pEC958 (brown).

### 3.5 The *pil* operon caused elevated protein interaction signals in pEK204

We examined the conjugative plasmid pEK204 further given the *pil* operon’s role in conjugation. 316 samples out of 1,060 with PPIs for both pEK204 and pEK499 had ≥1 of pEK204’s 14 *pilI-pilV* genes: the most common were *pilQ* (encoding a T4P secretin) and *pilT* (encoding a T4P twitching motility ATPase that helps retract the pilus). We scaled the Jaccard index for pEK204 by that for pEK499 to create a log2-scaled ratio that was positively correlated with the number of interacting host proteins for pEK204. This ratio identified samples with a higher potential for pEK204-specific PPIs, including several *Pseudomonas* samples (*P. aeruginosa, P. fluorescens* F113 as above) (Figure 5, Table S6). Some samples had >11 *pil* operon genes, including *Cellvibrio sp. BR*, whose 13 pEK204 proteins (Figure S17) had the highest number of pEK204-related PPIs of all samples (1,383, an excess of 7.4%), interacting with 660 other *Cellvibrio sp. BR* proteins (Figure S18). Samples with the highest pEK204-linked PPIs were associated with urine or water environments based on environmental niche information from BacDive [85], such as *Aromatoleum aromaticum* EbN1, a Betaproteobacterium from a river source. Similarly, water-associated *Rubrivivax gelatinosus* IL144 had 14 pEK204-matching proteins (Figure S19) that had PPIs with 507 other *Rubrivivax gelatinosus* IL144 proteins with 1,127 PPIs (an excess of 2%, Figure S20).

**Fig 5.**
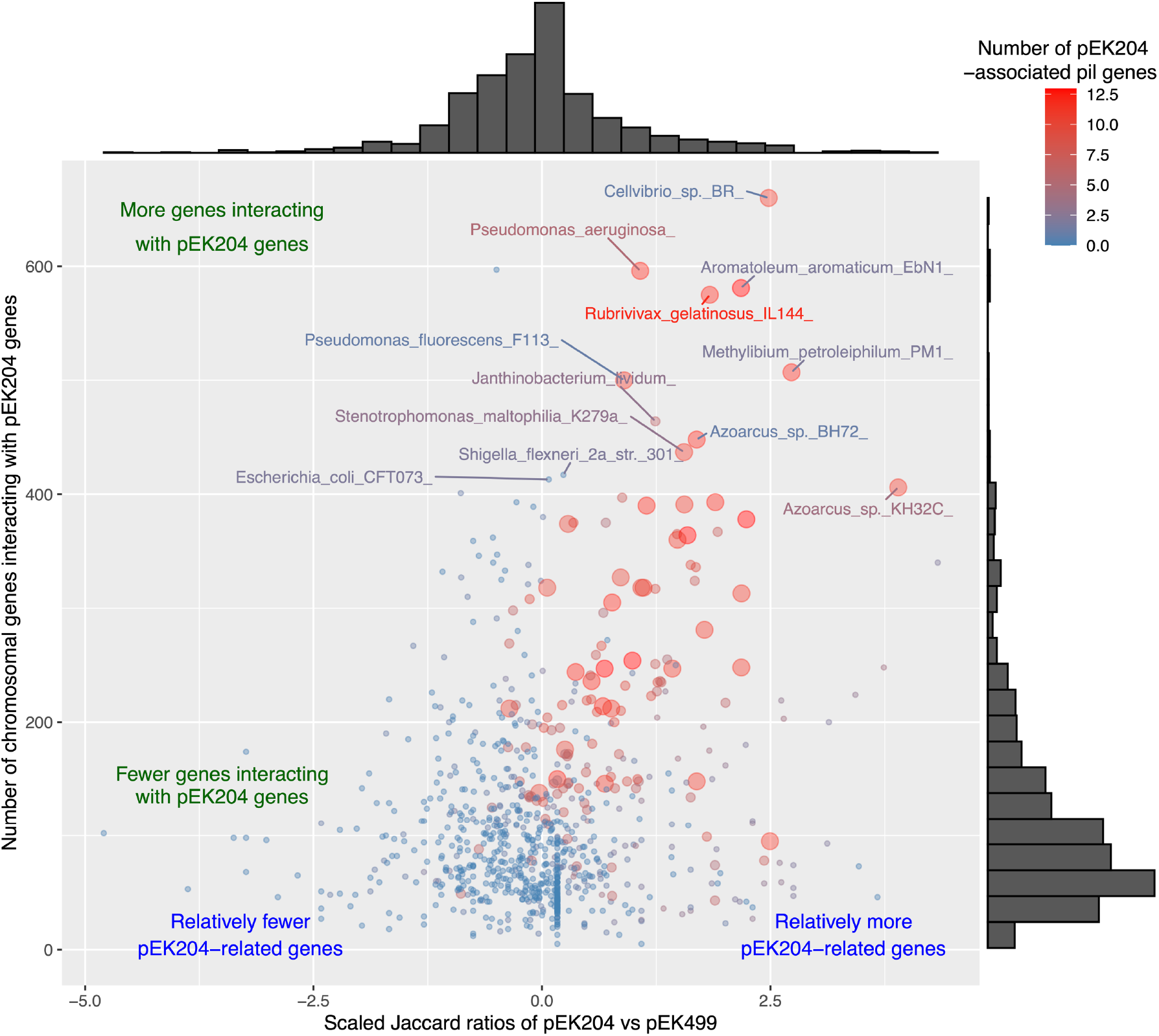
A plot of the normalised log2-scaled ratio of the Jaccard indices of pEK204 divided by that for pEK499 (x-axis) versus the number of sample proteins interacting with pEK204 proteins (y-axis). The data plotted is for 1,060 bacterial samples, of which 316 had ≥1 pEK204-linked *pil* gene(s) (14 genes, *pilIJKLMNOPQRSTUV*. The 12 labelled samples had ≥400 interacting host proteins and a log2-scaled ratio for pEK204 >0. A larger point size and a redder colour signifies the presence of more pEK204-related *pil* genes: blue indicates fewer genes. The histograms indicate the marginal densities of the scatterplots for both axes.

### 3.6 Interactions of *pil* operon gene products with other fimbrial adhesin proteins

The pairwise correlations of the 14 pEK204 *pil* genes’ frequencies across the 4,363 bacterial samples (Table S7) showed that most were positively or negatively correlated with one another, with the exceptions of *pilL, pilQ* and *pilV*. Previous work has shown that PilK-PilV, but not PilI or PilJ, are essential for R64 pilus generation [86]. Adding four *pap* genes and 14 *fim* genes to this identified five sets of genes, of which three involved *pil* genes. The first two sets were negatively correlated within their own groups, but were positively correlated with the other set (*pilO/P/R/T* and *pilA/I/J/M/N/S/U/W*/V). The 3^rd^ gene set was largely uncorrelated and stemmed from from FimT’s (a T4P biogenesis protein) and FimV’s (a motility protein) interactions with PilQ (a ComQ homolog), PilL and PilV (for FimT) and PilF/G/J (for FimV) in *P. aeruginosa* PAO1 and *Acinetobacter baumannii* A118 [87]. Testing for PPIs between *pil, fim* and *pap* proteins in the 4,363 samples found that *Ralstonia eutropha* H16 was the single sample with a Fim-Pil PPI across its 13 *pil* genes and four *fim* genes (though major structural protein FimA with PilM-O and pilQ, Figure S22A). Notably, both *E. coli* K12 MG1655 (Figure S22B) CFT073 (Figure S22C) had Fim and Pap PPIs, suggesting scope for Pil proteins to interact or evolve such PPIs.

## 4. Discussion

To explore the hypothesis that an elevated rate of plasmid-host PPIs might be associated with higher compatibility between a plasmid and a host cell, we tested for an excess of host-plasmid PPIs using eight key plasmids in representatives from most bacteria (n=4,363). We achieved this by modelling data from StringDB. We saw that these plasmids had extensive heterogeneity in their PPI levels across 24 *Enterobacterales* (*Escherichia, Klebsiella, Serratia* and *Shigella*) samples. This absence of a genus-level association was consistent with previous work [80].

To illuminate this issue, we explored the four genetically distinct *bla*OXA-48-positive plasmids in a larger set of 4,363 samples and found that these plasmids had ≥1 PPI with most bacteria, allowing us to identify instances where there was a large excess of PPIs in 40 (0.6%) sample-plasmid pairs. This identified *P. aeruginosa, A. aromaticum* EbN1 and 21 other common animal and plant pathogens as more likely to have PPIs with ≥1 of these *bla*OXA-48-positive plasmids, particularly with the pKP112 and pOXA-48 isoforms, suggesting those plasmids could become common in these species. Consistent with these predictions, *bla*OXA-48-positive human clinical isolates from Bangladesh [88] and Sudan [89] have been found in *P. aeruginosa*, even though previous *in vitro* experiments suggested incompatibility [34]. This is concerning because *bla*OXA-48 is the most common AMR gene in CPE and is prevalent in *Pseudomonas* based on carbapenemase gene rates in England from 2020-22 [15]. Given their already frequent retention of these plasmids, the high levels of carbapenems in clinical environments may allow these hosts to obtain, co-evolve with and compensate for the metabolic cost of possessing these plasmids [90].

We used the same approach to assess the other group of four genetically related plasmids (pEK204, pEK499, pEK516, pEC958) that likely originated in ExPEC [37]. ExPEC pose a significant public health problem and change their plasmid composition to improve their fitness [91]. Like the pOXA-48-related plasmids, these plasmids had ≥1 PPI with many bacteria, and we found large excess rates of PPIs in 48 species. 19% of these were from the *Pseudomonas* genus, where *P. fluorescens* F113 and *P. aeruginosa* had strong associations with pEK204. This was concerning because pEK204 has spread across pathogenic bacteria [26], and IncI1 plasmids like pEK204 promote AMR due to the presence of *bla*CTX-M, *bla*TEM, *bla*CMY and other ESBL genes [92-93].

*P. fluorescens* F113 and *P. aeruginosa* were among 12 samples with high rates of PPIs with pEK204, particularly in samples whose proteins interacted with *pil* operon proteins. This operon encodes a T4P with two main known isoforms [94], had 14 genes on pEK204, and is a virulence factor for *P. aeruginosa* [95-96]. A ten-gene *pil* operon is involved in conjugation of pathogenicity island PAPI-1 in *P. aeruginosa* PA14 independently of pEK204 [95] that has been linked to *P. aeruginosa* ST309 infection outbreaks [97], and *pil* genes form part of an integrative conjugative element Pac_ICE1 in *P. syringae* pv actinidiae [98]. Notably, *pil* may be more associated with aqueous environments, and the loss of *pil* genes from PAPI-1 has been associated with better acute human lung [99] and mouse gut [100] infection, perhaps due to Pil proteins acting as cell surface antigens [101]. Consistent with this connection to water, our approach identified bacteria associated with rivers like *Cellvibrio sp. BR* and others found in wastewater, sludge, sewage and freshwater *Rubrivivax gelatinosus* IL144 [100]. We identified three distinct groups of correlated *pil* gene rates in these 4,363 samples, suggesting that pEK204’s *pil* repertoire differed from other *pil* operon compositions. We found evidence of PPIs between *pil, fim* and *pap* gene products in *E. coli* and *R. eutropha*, which is important because ExPEC can possess type 1, IV and P fimbriae and tend to express a single fimbrial type at any time depending on the host niche, indicating coordinated expression [103-105]. In this way, the gain of *pil* via pEK204 could facilitate superior host survival, conjugation or biofilm formation in water environments [41].

A limitation of this study was the inconsistent nomenclature of plasmid and samples, which meant that certain protein matches were not detected or misclassified, and affecting PPI identification. For instance, functions were assigned to ∼46% of genes in previous work [106]. In addition, we did not map reads or align sequences here to detect genes and relied on the accuracy of existing gene annotation tools based on protein sequence patterns. Such DNA sequence analysis would yield diverse and likely ambiguous gene cluster alignment scores that also lead to matches being unclear or imprecisely categorised. Future work is needed to improve genome and plasmid annotation, explore the strength and specificity of PPIs, and assess other highly transmissible plasmids.

## Conclusions

Given that no study yet has explored PPI between key plasmids across all bacteria to date, so we developed a model testing for excess rates of PPIs between 4,363 bacterial proteomes and the proteomes of eight plasmids of public health concern. 23 potential bacterial host cells had high PPI rates with four *bla*OXA-48-positive plasmids, including *P. aeruginosa*. Similarly, 48 species had elevated PPI rates with four common *E. coli* plasmids, including nine from the *Pseudomonas* genus. One of the latter plasmids (pEK204), is associated with host cell binding and conjugative transfer in aqueous environments, and appeared to be a major driver of host-plasmid PPIs in the species with high rates of PPIs with pEK204. These results overcome a limitation that the gain of a new plasmid may have heterogenous effects on different host cells, and this can depend on genetic differentiation levels, as has been illustrated for pOXA-48 [80]. Consequently, our approach using excess rates of PPIs between plasmid and host cell proteins added results on the potential compatibility of plasmids with particular species. Our study has implications for controlling emerging problems, such as *bla*OXA-48-carrying plasmids that can spread among numerous diverse host genera.

## ^1^ Abbreviations

(AMR): Antimicrobial resistance
(CPE): Carbapenemase-producing Enterobacteriaceae
(ExPEC): Extraintestinal pathogenic E. coli
(HGT): Horizontal gene transfer
(PPIs): Protein-protein interactions
(T4P): Type IV pilus

## Author Contributions

TD – Conceptualization, Methodology, Software, Formal analysis, Investigation, Data Curation, Writing - Original Draft, Writing - Review & Editing, Visualisation, Project administration. MJL – Methodology, Formal analysis, Investigation, Data Curation, Visualisation. CA – Formal analysis, Investigation, Data Curation, Visualisation. AMcD – Formal analysis, Investigation, Data Curation, Visualisation. AR – Conceptualization, Methodology, Writing - Review & Editing.

## Acknowledgements

The authors acknowledge School of Biotechnology (Dublin City University) student funding. We thanks the reviewers and others with comments for their constructive input.

## Data Summary

Processing files and relevant code associated with this paper are available on Github at https://github.com/downingtim/plasmids_2022.

S. Kurtz, A. Phillippy, A.L. Delcher, M. Smoot, M. Shumway, C. Antonescu, S. L. Salzberg, Versatile and open software for comparing large genomes, Genome Biol. 5 (2004) 1–9.

