## Supplementary material for "Informing plasmid compatibility with bacterial hosts using protein-protein interaction data": Suppl_Data

Supplementary Data

**Page	Item	Description**

1	Table S1	The PPI data for four pOXA-48 plasmid isoforms across 24 *Enterobacterales*.

2	Table S2	The PPI data for four *Enterobacterales*-related plasmids across 24

*Enterobacterales.*

3	Table S3	Details of the linear models predicting the number of PPIs per sample-plasmid pair.

4	Table S4	Key data sample-plasmid pairs for pOXA-48-related isoforms.

5	Table S5	Key data sample-plasmid pairs for pEK204, pEK499, pEK516 and pEC958.

6	Table S6	The frequencies of the 14 pEK204-related pil genes in bacterial samples.

7	Table S7	The pairwise PPIs between *fim* and *pap* genes observed for *E. coli.*

8	Figure S1	The numbers of interacting host proteins per sample and PPIs per sample for the

pOXA-48- related isoforms.

9	Figure S2	The numbers of PPIs per sample relative to the numbers of interacting host proteins

per sample for the four pOXA-48 isoforms.

10	Figure S3	The residuals of PPIs with each plasmid per sample versus by the number of host

proteins interacting with each plasmid for the four pOXA-48 isoforms.

11	Figure S4	A PPI network for *Pseudomonas aeruginosa*’s proteins on pKP112.

11	Figure S5	A PPI network showing pKP112-related PPIs interacting with *Pseudomonas*

*aeruginosa*’s proteins.

12	Figure S6	A PPI network for *Aromatoleum* *aromaticum* EbN1’s proteins on pOXA-48.

12	Figure S7	A PPI network showing pOXA-48-related PPIs interacting with *Aromatoleum*

*aromaticum* EbN1.

13	Figure S8	A PPI network for *Enterobacter cloacae subsp. cloacae* ATCC13047‘s proteins on

pKP112.

13	Figure S9	A PPI network showing pKP112-related PPIs interacting with *Enterobacter*

*cloacae subsp. cloacae* ATCC13047.

14	Figure S10	The numbers of interacting host proteins per sample and PPIs per sample for

pEC958, pEK204, pEK499 and pEK516.

15	Figure S11	The numbers of PPIs per sample relative to the numbers of interacting host proteins

per sample for pEC958, pEK204, pEK499 and pEK516.

16	Figure S12	The residuals of PPIs with each plasmid per sample versus by the number of host

proteins interacting with each plasmid for pEC958, pEK204, pEK499 and pEK516.

17	Figure S13	A PPI network for *Pseudomonas aeruginosa*’s proteins on pEK204.

17	Figure S14	A PPI network showing pEK204-related PPIs interacting with *Pseudomonas*

*aeruginosa*’s proteins.

18	Figure S15	A PPI network for *Aromatoleum* *aromaticum* EbN1’s proteins on pEK204.

18	Figure S16	A PPI network showing pEK204-related PPIs interacting with *Aromatoleum*

*aromaticum* EbN1’s proteins.

19	Figure S17	A PPI network for *Cellvibrio sp. BR*'s proteins on pEK204.

19	Figure S18	A PPI network showing pEK204-related PPIs interacting with *Cellvibrio sp. BR*'s

Proteins.

20	Figure S19	A PPI network for *Rubrivivax gelatinosus* IL144’s proteins on pEK204.

20	Figure S20	A PPI network showing pEK204-related PPIs interacting with *Rubrivivax*

*gelatinosus* IL144’s proteins.

21	Figure S21	The pairwise association based on hierarchical clustering between known *fim, pap*

and *pil* genes in bacteria.

22	Figure S22	PPI networks for (A) *Ralstonia eutropha* H16, (B) *E. coli* K12 MG1655 and (C) *E.*

*coli* CFT073.

| Plasmid name | pOXA-48 | | | pKP112 | | | pOXA-48_30715 | | | pOXA-48_k8 | | |
| --- | --- | --- | --- | --- | --- | --- | --- | --- | --- | --- | --- | --- |
| *Klebsiella pneumoniae* strain | Kp11978 | | | KP112 | | | Kpn-30715/15 | | | K8 | | |
| Accession number | JN626286.1 | | | LN864819.1 | | | NZ_KX523901.1 | | | MT441554.1 | | |
| Length (bp) | 61,881 | | | 84,252 | | | 65,488 | | | 65,499 | | |
| Number of genes | 40 | | | 62 | | | 84 | | | 87 | | |
| Number of unique genes | 37 | | | 57 | | | 12 | | | 18 | | |
| Number of CDS elements | 74 | | | 102 | | | 84 | | | 87 | | |
| **Sample** | **Jaccard** | **PPIs** | **Genes** | **Jaccard** | **PPIs** | **Genes** | **Jaccard** | **PPIs** | **Genes** | **Jaccard** | **PPIs** | **Genes** |
| *Escherichia albertii* KF1 | 0.028 | 125 | 144 | 0.025 | 114 | 134 | 0.017 | 77 | 76 | 0.021 | 95 | 97 |
| *Escherichia albertii* TW07627 | 0.009 | 58 | 41 | 0.007 | 49 | 31 | 0.000 | 0 | 0 | 0.009 | 37 | 38 |
| *Escherichia coli* 536 | 0.009 | 63 | 44 | 0.009 | 63 | 44 | 0.004 | 19 | 20 | 0.008 | 37 | 37 |
| *Escherichia coli* ATCC 8739 | 0.010 | 60 | 43 | 0.012 | 65 | 49 | 0.005 | 22 | 23 | 0.012 | 54 | 52 |
| *Escherichia coli* BL21 | 0.030 | 139 | 126 | 0.022 | 106 | 92 | 0.017 | 84 | 73 | 0.028 | 130 | 116 |
| *Escherichia coli* CFT073 | 0.019 | 117 | 103 | 0.019 | 116 | 102 | 0.013 | 79 | 71 | 0.021 | 123 | 113 |
| *Escherichia coli* O157:H7 str. EDL933 | 0.017 | 95 | 99 | 0.011 | 64 | 67 | 0.004 | 20 | 21 | 0.020 | 113 | 116 |
| *Escherichia coli* str. K-12 substr. MG1655 | 0.037 | 173 | 154 | 0.029 | 142 | 122 | 0.028 | 148 | 117 | 0.047 | 239 | 195 |
| *Escherichia hermannii* NBRC 105704 | 0.013 | 72 | 54 | 0.011 | 66 | 47 | 0.010 | 54 | 42 | 0.019 | 92 | 80 |
| *Escherichia vulneris* NBRC 102420 | 0.031 | 154 | 133 | 0.029 | 144 | 122 | 0.020 | 83 | 84 | 0.036 | 156 | 150 |
| *Shigella flexneri* 2a str. 301 | 0.033 | 180 | 147 | 0.029 | 164 | 130 | 0.019 | 94 | 84 | 0.033 | 166 | 148 |
| *Klebsiella oxytoca* | 0.023 | 152 | 135 | 0.030 | 221 | 173 | 0.009 | 50 | 51 | 0.014 | 82 | 82 |
| *Klebsiella pneumoniae* | 0.008 | 45 | 46 | 0.008 | 45 | 46 | 0.000 | 0 | 0 | 0.005 | 28 | 29 |
| *Serratia fonticola* AU-AP2C | 0.000 | 0 | 0 | 0.000 | 0 | 0 | 0.000 | 0 | 0 | 0.003 | 13 | 12 |
| *Serratia fonticola* RB-25 | 0.007 | 34 | 33 | 0.007 | 34 | 33 | 0.000 | 0 | 0 | 0.005 | 23 | 22 |
| *Serratia g*rimesii | 0.007 | 29 | 30 | 0.007 | 29 | 30 | 0.000 | 0 | 0 | 0.007 | 29 | 30 |
| *Serratia marcescens* FGI94 | 0.007 | 33 | 32 | 0.007 | 33 | 32 | 0.000 | 0 | 0 | 0.004 | 18 | 17 |
| *Serratia marcescens* subsp. marcescens Db11 | 0.024 | 131 | 115 | 0.020 | 114 | 97 | 0.016 | 78 | 76 | 0.020 | 101 | 93 |
| *Serratia plymuthica* | 0.007 | 32 | 33 | 0.007 | 32 | 33 | 0.000 | 0 | 0 | 0.004 | 17 | 18 |
| *Serratia proteamaculans* 568 | 0.006 | 28 | 29 | 0.006 | 28 | 29 | 0.000 | 0 | 0 | 0.003 | 16 | 17 |
| *Serratia* sp. Ag1 | 0.008 | 36 | 35 | 0.008 | 36 | 35 | 0.000 | 0 | 0 | 0.005 | 23 | 22 |
| *Serratia* sp. ATCC 39006 | 0.008 | 33 | 34 | 0.008 | 33 | 34 | 0.000 | 0 | 0 | 0.002 | 10 | 11 |
| *Serratia* sp. DD3 | 0.030 | 140 | 155 | 0.064 | 306 | 333 | 0.000 | 0 | 0 | 0.032 | 151 | 150 |
| *Serratia* sp. M24T3 | 0.005 | 22 | 23 | 0.005 | 22 | 23 | 0.000 | 0 | 0 | 0.005 | 22 | 23 |

**Table S1.** The PPI data for the four main pOXA-48 plasmid isoforms across 24 samples from three genera: *Escherichia*/*Shigella* (n=11), *Klebsiella* (n=2) and *Serratia* (n=11). The interaction data is represented as the Jaccard index of the fraction of interacting genes relative to the combined total of sample and plasmid genes per combination, then the total numbers of PPIs, and the number of interacting genes per sample.

| Plasmid name | pEK204 | | | | pEK499 | | | | pEK516 | | | | pEC958 | | |
| --- | --- | --- | --- | --- | --- | --- | --- | --- | --- | --- | --- | --- | --- | --- | --- |
| *Escherichia coli* strain | C | | | | A | | | | D | | | | EC958 | | |
| Accession number | EU935740 | | | | EU935739 | | | | EU935738 | | | | NZ_HG941719.1 | | |
| Length (bp) | 93,732 | | | | 117,536 | | | | 64,471 | | | | 135,602 | | |
| Number of genes | 89 | | | | 103 | | | | 65 | | | | 153 | | |
| Number of unique genes | 86 | | | | 84 | | | | 55 | | | | 42 | | |
| Number of CDS elements | 95 | | | | 141 | | | | 79 | | | | 153 | | |
| **Sample** | **Jaccard** | **PPIs** | **Genes** | **Jaccard** | | **PPIs** | **Genes** | **Jaccard** | | **PPIs** | **Genes** | **Jaccard** | | **PPIs** | **Genes** |
| *Escherichia a*lbertii KF1 | 0.026 | 127 | 117 | 0.041 | | 198 | 183 | 0.017 | | 76 | 77 | 0.017 | | 76 | 77 |
| *Escherichia albertii* TW07627 | 0.010 | 63 | 44 | 0.029 | | 169 | 130 | 0 | | 0 | 0 | 0 | | 0 | 0 |
| *Escherichia coli* 536 | 0.030 | 148 | 140 | 0.042 | | 225 | 199 | 0.029 | | 135 | 137 | 0.017 | | 77 | 80 |
| *Escherichia coli* ATCC 8739 | 0.033 | 153 | 142 | 0.046 | | 213 | 197 | 0.03 | | 123 | 127 | 0.015 | | 62 | 64 |
| *Escherichia coli* BL21 | 0.050 | 222 | 215 | 0.08 | | 374 | 343 | 0.082 | | 351 | 347 | 0.038 | | 157 | 161 |
| *Escherichia coli* CFT073 | 0.065 | 373 | 355 | 0.069 | | 406 | 376 | 0.076 | | 418 | 413 | 0.022 | | 115 | 118 |
| *Escherichia coli* O157:H7 str. EDL933 | 0.038 | 221 | 222 | 0.057 | | 389 | 333 | 0.027 | | 157 | 158 | 0.032 | | 192 | 185 |
| *Escherichia coli* str. K-12 substr. MG1655 | 0.091 | 406 | 384 | 0.142 | | 684 | 597 | 0.113 | | 478 | 474 | 0.057 | | 233 | 236 |
| *Escherichia hermannii* NBRC 105704 | 0.032 | 146 | 135 | 0.078 | | 381 | 332 | 0.029 | | 118 | 122 | 0.023 | | 95 | 96 |
| *Escherichia vulneris* NBRC 102420 | 0.044 | 184 | 189 | 0.084 | | 420 | 359 | 0.048 | | 201 | 204 | 0.046 | | 193 | 196 |
| *Shigella flexneri* 2a str. 301 | 0.092 | 458 | 417 | 0.088 | | 438 | 398 | 0.059 | | 273 | 266 | 0.041 | | 202 | 184 |
| *Klebsiella oxytoca* | 0.032 | 187 | 190 | 0.059 | | 381 | 346 | 0.04 | | 245 | 236 | 0.028 | | 159 | 165 |
| *Klebsiella pneumoniae* | 0 | 0 | 0 | 0.008 | | 46 | 48 | 0 | | 0 | 0 | 0 | | 0 | 0 |
| *Serratia fonticola* AU-AP2C | 0 | 0 | 0 | 0.008 | | 32 | 35 | 0 | | 0 | 0 | 0.001 | | 4 | 5 |
| *Serratia fonticola* RB-25 | 0 | 0 | 0 | 0.004 | | 19 | 20 | 0 | | 0 | 0 | 0 | | 0 | 0 |
| *Serratia g*rimesii | 0 | 0 | 0 | 0.01 | | 45 | 47 | 0 | | 0 | 0 | 0 | | 0 | 0 |
| *Serratia marcescens* FGI94 | 0 | 0 | 0 | 0.004 | | 16 | 17 | 0 | | 0 | 0 | 0 | | 0 | 0 |
| *Serratia marcescens* subsp. marcescens Db11 | 0.017 | 84 | 84 | 0.056 | | 283 | 267 | 0.026 | | 123 | 126 | 0.034 | | 158 | 162 |
| *Serratia plymuthica* | 0 | 0 | 0 | 0.007 | | 34 | 36 | 0 | | 0 | 0 | 0 | | 0 | 0 |
| *Serratia proteamaculans* 568 | 0 | 0 | 0 | 0.007 | | 35 | 37 | 0 | | 0 | 0 | 0 | | 0 | 0 |
| *Serratia* sp. Ag1 | 0 | 0 | 0 | 0.005 | | 22 | 24 | 0 | | 0 | 0 | 0.001 | | 4 | 5 |
| *Serratia* sp. ATCC 39006 | 0.009 | 39 | 41 | 0.007 | | 32 | 33 | 0 | | 0 | 0 | 0.013 | | 55 | 56 |
| *Serratia* sp. DD3 | 0.033 | 169 | 158 | 0.05 | | 235 | 238 | 0.032 | | 153 | 154 | 0.026 | | 120 | 121 |
| *Serratia* sp. M24T3 | 0 | 0 | 0 | 0 | | 0 | 0 | 0 | | 0 | 0 | 0 | | 0 | 0 |

**Table S2.** The PPI data for pEK204, pEK499, pEK516 and pEC958 across the same 24 samples from three genera: *Escherichia*/*Shigella* (n=11), *Klebsiella* (n=2) and *Serratia* (n=11). The interaction data is represented as the Jaccard index of the fraction of interacting genes relative to the combined total of sample and plasmid genes per combination, then the total numbers of PPIs, and the number of interacting genes per sample.

| **Dataset** | | | | **All pairs** | | **pOXA-48** | | **pKP112** | | **pOXA-48_30715** | | **pOXA-48_k8** | |
| --- | --- | --- | --- | --- | --- | --- | --- | --- | --- | --- | --- | --- | --- |
| Number of pairs | | | | 6,181 | | 1,820 | | 1,868 | | 673 | | 1,820 | |
| Baseline number of PPIs per sample p value | | | | 1.9e-14 | | 3.7e-10 | | 6.4e-7 | | 0.001 | | 2.3e-15 | |
| Number of interacting proteins | r | | | 0.98 | | 0.96 | | 0.96 | | 0.96 | | 0.995 | |
|  | p value | | | 4.6e-15 | | 4.6e-15 | | 4.6e-15 | | 4.6e-15 | | 2.3e-15 | |
| Number of proteins per sample | r | | | <0.01 | | 0.11 | | 0.13 | | 0.38 | | 0.085 | |
|  | p value | | | >0.5 | | >0.5 | | >0.5 | | >0.5 | | 0.064 | |
| Interaction of proteins per sample with interacting proteins p value | | | | >0.5 | | >0.5 | | >0.5 | | >0.5 | | 0.014 | |
| **Dataset** | | | | | **All pairs** | | **pEK204** | | **pEK499** | | **pEK516** | | **pEC958** |
| Number of pairs | | | | | 3,828 | | 1,820 | | 1,868 | | 673 | | 1,820 |
| Baseline number of PPIs per sample p value | | | | | 0.5 | | 1.4e-6 | | 8.3e-8 | | 0.026 | | 0.26 |
| Number of interacting genes | | r | 0.89 | | | | 0.92 | | 0.98 | | 0.96 | | 0.84 |
|  |  | p value | 4.6e-15 | | | | 4.6e-15 | | 4.6e-15 | | 4.6e-15 | | 4.6e-15 |
| Number of proteins per sample | | r | 0.25 | | | | 0.34 | | 0.18 | | 0.32 | | 0.30 |
|  |  | p value | >0.5 | | | | >0.5 | | >0.5 | | >0.5 | | >0.5 |
| Interaction of proteins per sample with interacting proteins p value | | | | | >0.5 | | >0.5 | | >0.5 | | >0.5 | | >0.5 |

**Table S3**. Output of the linear models predicting the number of PPIs per sample-plasmid pair based on the numbers of interacting host proteins, plasmid proteins, total proteins in the sample, and associated pairwise interactions. The upper data is shown for all pOXA-48-related isoforms, pOXA-48, pKP112, pOXA-48_30715 and pOXA-48_k8. The lower data is for pEC958, pEK204, pEK499 and pEK516. The correlation coefficients were based on the linear correlation of the variable with the number of interactions. The number of interacting proteins and the baseline number of PPIs per sample were useful predictors, whereas the total number of proteins per sample and its interaction with the number of interacting proteins were not – except for pOXA-48_k8 and pEC958 where the latter two variable had slight effects.

| **Plasmid** | **Sample** | **Number of interacting host proteins** | **Num of PPIs** | **Expected PPIs** | **Excess Fraction of PPIs** | **Total sample proteins** | **Total sample PPIs** |
| --- | --- | --- | --- | --- | --- | --- | --- |
| pKP112 | *Pseudomonas aeruginosa* | 450 | 684 | 539.7 | 0.267 | 6,299 | 131,860 |
| pKP112 | *Aromatoleum aromaticum EbN1* | 297 | 452 | 354.2 | 0.276 | 4,359 | 54,096 |
| pOXA-48 | *Aromatoleum aromaticum EbN1* | 252 | 400 | 299.7 | 0.335 | 4,359 | 54,096 |
| pKP112 | *Ralstonia solanacearum GMI1000* | 249 | 358 | 295.8 | 0.210 | 5,778 | 78,200 |
| pOXA-48 | *Ralstonia solanacearum GMI1000* | 249 | 358 | 295.7 | 0.211 | 5,778 | 78,200 |
| pOXA-48 | *Buttiauxella agrestis ATCC 33320* | 211 | 336 | 250.0 | 0.344 | 4,331 | 86,662 |
| pKP112 | *Glaciecola lipolytica E3* | 203 | 356 | 240.1 | 0.483 | 4,331 | 67,056 |
| pKP112 | *Buttiauxella agrestis ATCC 33320* | 195 | 320 | 230.4 | 0.389 | 4,331 | 86,662 |
| pOXA-48_k8 | *Methyloglobulus morosus KoM1* | 183 | 233 | 193.6 | 0.203 | 3,737 | 53,749 |
| pKP112 | *Prevotella buccae ATCC 33574* | 170 | 253 | 200.3 | 0.263 | 2,896 | 35,597 |
| pKP112 | *Legionella oakridgensis ATCC 33761 DSM 21215* | 166 | 353 | 195.4 | 0.807 | 2,944 | 29,151 |
| pOXA-48 | *Legionella oakridgensis ATCC 33761 DSM 21215* | 166 | 353 | 195.8 | 0.803 | 2,944 | 29,151 |
| pKP112 | *Bacteroides fragilis NCTC 9343* | 157 | 523 | 184.3 | 1.838 | 4,236 | 58,209 |
| pOXA-48 | *Bacteroides fragilis NCTC 9343* | 151 | 503 | 177.2 | 1.838 | 4,236 | 58,209 |
| pKP112 | *Methyloglobulus morosus KoM1* | 148 | 243 | 173.4 | 0.401 | 3,737 | 53,749 |
| pKP112 | *Achromobacter xylosoxidans A8* | 147 | 228 | 171.8 | 0.327 | 6,717 | 115,251 |
| pKP112 | *Lactobacillus plantarum WCFS1* | 146 | 228 | 171.1 | 0.333 | 3,109 | 36,185 |
| pKP112 | *Candidatus Glomeribacter gigasporarum BEG34* | 143 | 211 | 167.7 | 0.258 | 1,485 | 19,518 |
| pOXA-48_k8 | *Lactobacillus plantarum WCFS1* | 141 | 182 | 148.6 | 0.225 | 3,109 | 36,185 |
| pKP112 | *Methylibium petroleiphilum PM1* | 138 | 208 | 161.2 | 0.290 | 4,517 | 60,002 |
| pOXA-48 | *Candidatus Glomeribacter gigasporarum BEG34* | 137 | 202 | 161.1 | 0.254 | 1,485 | 19,518 |
| pKP112 | *Acidovorax sp. KKS102* | 135 | 323 | 157.5 | 1.051 | 4,758 | 57,517 |
| pOXA-48 | *Lactobacillus plantarum WCFS1* | 135 | 217 | 158.2 | 0.372 | 3,109 | 36,185 |
| pKP112 | *Chryseobacterium indologenes NBRC 14944* | 134 | 231 | 156.4 | 0.477 | 4,192 | 43,715 |
| pOXA-48 | *Methyloglobulus morosus KoM1* | 128 | 208 | 149.5 | 0.392 | 3,737 | 53,749 |
| pKP112 | *Enterobacter cloacae subsp. cloacae ATCC 13047* | 124 | 173 | 144.1 | 0.201 | 5,414 | 79,364 |
| pKP112 | *Legionella longbeachae NSW150* | 122 | 176 | 141.9 | 0.240 | 3,478 | 38,062 |
| pOXA-48 | *Legionella longbeachae NSW150* | 122 | 176 | 142.3 | 0.237 | 3,478 | 38,062 |
| pKP112 | *Burkholderia multivorans ATCC 17616* | 121 | 186 | 140.3 | 0.326 | 6,262 | 73,063 |
| pKP112 | *Rickettsia bellii RML369-C* | 120 | 199 | 139.7 | 0.424 | 1,434 | 18,297 |
| pOXA-48 | *Rickettsia bellii RML369-C* | 120 | 199 | 140.5 | 0.417 | 1,434 | 18,297 |
| pOXA-48_30715 | *Prevotella sp. oral taxon 472 str. F0295* | 114 | 183 | 120.6 | 0.517 | 3,092 | 36,501 |
| pOXA-48 | *Citrobacter farmeri GTC 1319* | 113 | 262 | 131.0 | 0.999 | 4,560 | 66,037 |
| pOXA-48_k8 | *Citrobacter farmeri GTC 1319* | 111 | 141 | 116.3 | 0.213 | 4,560 | 66,037 |
| pKP112 | *Citrobacter farmeri GTC 1319* | 109 | 259 | 126.0 | 1.056 | 4,560 | 66,037 |
| pOXA-48 | *Methylibium petroleiphilum PM1* | 108 | 168 | 125.0 | 0.344 | 4,517 | 60,002 |
| pKP112 | *Helicobacter hepaticus ATCC 51449* | 106 | 148 | 122.7 | 0.206 | 1,877 | 23,481 |
| pKP112 | *Cupriavidus metallidurans CH34* | 105 | 152 | 120.9 | 0.257 | 6,169 | 90,771 |
| pOXA-48 | *Chryseobacterium indologenes NBRC 14944* | 104 | 200 | 120.2 | 0.664 | 4,192 | 43,715 |
| pOXA-48_30715 | *Bacteroides fragilis NCTC 9343* | 101 | 193 | 105.9 | 0.822 | 4,236 | 58,209 |

**Table S4**. Key data sample-plasmid pairs with a high excess fraction of PPIs (>0.2) and >100 interacting sample proteins and >100 PPIs between the plasmid proteins and the other sample proteins for the four pOXA-48-related isoforms, pOXA-48 (37 unique proteins), pKP112 (57 unique proteins), pOXA-48_30715 (12 unique proteins) and pOXA-48_k8 (18 unique proteins). The table is sorted by the number of interacting host proteins.

| **Plasmid** | **Sample** | **Number of interacting host proteins** | **Num of PPIs** | **Expected PPIs** | **Excess Fraction of PPIs** | **Total sample proteins** | **Total sample PPIs** |
| --- | --- | --- | --- | --- | --- | --- | --- |
| pEK204 | *Pseudomonas fluorescens F113* | 500 | 1,215 | 951.0 | 5,862 | 155,946 | 0.217 |
| pEK204 | *Azoarcus sp. KH32C* | 406 | 1,049 | 764.5 | 5,192 | 104,270 | 0.271 |
| pEK204 | *Pseudomonas sp. BAY1663* | 364 | 1,120 | 680.5 | 5,056 | 124,944 | 0.392 |
| pEK204 | *Candidatus Symbiobacter mobilis CR* | 336 | 829 | 633.7 | 2,626 | 53,905 | 0.236 |
| pEK204 | *Ralstonia solanacearum GMI1000* | 327 | 847 | 603.2 | 5,778 | 78,200 | 0.288 |
| pEK204 | *Burkholderia pseudomallei K96243* | 324 | 790 | 597.3 | 5,736 | 101,044 | 0.244 |
| pEK204 | *Pseudomonas knackmussii B13* | 318 | 735 | 584.8 | 5,837 | 91,098 | 0.204 |
| pEK204 | *Marinobacter sp. HL-58* | 305 | 717 | 567.2 | 3,691 | 73,964 | 0.209 |
| pEC958 | *Buttiauxella agrestis ATCC 33320* | 291 | 514 | 372.9 | 4,331 | 86,662 | 0.275 |
| pEK204 | *Pseudomonas sp. 20 BN* | 254 | 874 | 465.6 | 3,414 | 80,385 | 0.467 |
| pEK204 | *Cupriavidus metallidurans CH34* | 248 | 668 | 442.6 | 6,169 | 90,771 | 0.337 |
| pEK204 | *endosymbiont of Tevnia jerichonana* | 247 | 618 | 452.2 | 3,230 | 58,445 | 0.268 |
| pEK204 | *Pseudomonas stutzeri A1501* | 247 | 648 | 448.6 | 4,144 | 65,185 | 0.308 |
| pEK204 | *Cupriavidus taiwanensis LMG 19424* | 245 | 636 | 437.7 | 5,897 | 94,092 | 0.312 |
| pEK204 | *Collimonas fungivorans Ter331* | 244 | 668 | 441.4 | 4,433 | 52,509 | 0.339 |
| pEK204 | *gamma proteobacterium HdN1* | 236 | 562 | 428.0 | 3,763 | 46,422 | 0.238 |
| pEK204 | *Legionella oakridgensis ATCC 33761 DSM 21215* | 236 | 767 | 431.2 | 2,944 | 29,151 | 0.438 |
| pEK204 | *Geobacter daltonii FRC-32* | 235 | 554 | 425.8 | 3,798 | 62,966 | 0.231 |
| pEK204 | *Pseudomonas resinovorans NBRC 106553* | 223 | 596 | 393.5 | 5,875 | 94,210 | 0.340 |
| pEK204 | *Citrobacter farmeri GTC 1319* | 222 | 565 | 396.7 | 4,560 | 66,037 | 0.298 |
| pEC958 | *Ralstonia solanacearum GMI1000* | 221 | 398 | 280.7 | 5,778 | 78,200 | 0.295 |
| pEK204 | *Chromobacterium violaceum ATCC 12472* | 219 | 511 | 391.2 | 4,414 | 72,864 | 0.234 |
| pEK204 | *Massilia sp. LC238* | 217 | 498 | 384.5 | 5,101 | 109,297 | 0.228 |
| pEK499 | *Erwinia tasmaniensis Et1-99* | 215 | 301 | 238.1 | 3,473 | 48,850 | 0.209 |
| pEK204 | *Pseudomonas pseudoalcaligenes CECT 5344* | 215 | 492 | 383.5 | 4,324 | 56,980 | 0.221 |
| pEK204 | *Xanthomonas axonopodis pv. citri str. 306* | 214 | 560 | 381.3 | 4,365 | 59,510 | 0.319 |
| pEK204 | *Ralstonia solanacearum PSI07* | 209 | 535 | 368.9 | 4,975 | 65,014 | 0.311 |
| pEK204 | *Pseudomonas syringae pv. phaseolicola 1448A* | 200 | 468 | 350.1 | 5,129 | 83,090 | 0.252 |
| pEK204 | *Glaciecola lipolytica E3* | 195 | 610 | 343.2 | 4,331 | 67,056 | 0.437 |
| pEK204 | *Shewanella oneidensis MR-1* | 187 | 425 | 328.1 | 4,086 | 70,578 | 0.228 |
| pEC958 | *Citrobacter farmeri GTC 1319* | 179 | 433 | 225.4 | 4,560 | 66,037 | 0.480 |
| pEC958 | *Legionella oakridgensis ATCC 33761 DSM 21215* | 179 | 509 | 225.3 | 2,944 | 29,151 | 0.557 |
| pEK499 | *Bacteroides fragilis NCTC 9343* | 170 | 442 | 187.1 | 4,236 | 58,209 | 0.577 |
| pEK204 | *Legionella longbeachae NSW150* | 162 | 392 | 280.2 | 3,478 | 38,062 | 0.285 |
| pEK204 | *Pseudoalteromonas spongiae UST010723-006* | 156 | 405 | 265.3 | 4,191 | 64,372 | 0.345 |
| pEK204 | *Alcanivorax borkumensis SK2* | 152 | 333 | 262.9 | 2,759 | 38,987 | 0.211 |
| pEK204 | *Acinetobacter calcoaceticus PHEA-2* | 150 | 408 | 255.5 | 3,603 | 42,697 | 0.374 |
| pEK204 | *Nitrosomonas europaea ATCC 19718* | 150 | 371 | 259.9 | 2,489 | 31,078 | 0.299 |
| pEK204 | *Geobacter sulfurreducens PCA* | 148 | 344 | 247.4 | 4,632 | 60,394 | 0.281 |
| pEK204 | *Janthinobacterium sp. Marseille* | 148 | 436 | 251.1 | 3,698 | 50,013 | 0.424 |
| pEK204 | *Ralstonia solanacearum Po82* | 147 | 339 | 250.5 | 3,338 | 34,934 | 0.261 |
| pEK204 | *Psychrobacter arcticus 273-4* | 146 | 354 | 253.3 | 2,130 | 30,388 | 0.284 |
| pEK516 | *Bacteroides fragilis NCTC 9343* | 144 | 417 | 151.8 | 4,236 | 58,209 | 0.636 |
| pEK204 | *Photorhabdus luminescens subsp. laumondii TTO1* | 144 | 312 | 239.0 | 4,742 | 65,744 | 0.234 |
| pEK204 | *Bacteroides fragilis NCTC 9343* | 143 | 773 | 238.9 | 4,236 | 58,209 | 0.691 |
| pEK204 | *Legionella pneumophila subsp. pneumophila str. Philadelphia 1* | 142 | 358 | 242.0 | 2,949 | 36,859 | 0.324 |
| pEK204 | *Ramlibacter tataouinensis TTB310* | 142 | 396 | 238.3 | 3,881 | 50,630 | 0.398 |
| pEK204 | *Vibrio vulnificus* | 134 | 332 | 219.7 | 4,512 | 77,958 | 0.338 |
| pEK204 | *Legionella shakespearei DSM 23087* | 129 | 325 | 215.9 | 2,931 | 38,506 | 0.336 |
| pEK204 | *Candidatus Hamiltonella defensa 5AT* | 120 | 401 | 200.8 | 2,185 | 23,667 | 0.499 |
| pEC958 | *Bacteroides fragilis NCTC 9343* | 118 | 485 | 145.0 | 4,236 | 58,209 | 0.701 |
| pEK204 | *Azotobacter vinelandii DJ* | 110 | 351 | 169.2 | 5,075 | 74,688 | 0.518 |
| pEK204 | *Vibrio azureus NBRC 104587* | 110 | 431 | 172.8 | 4,158 | 68,901 | 0.599 |
| pEK204 | *Prevotella nigrescens ATCC 33563* | 109 | 443 | 177.7 | 2,421 | 28,879 | 0.599 |
| pEK204 | *Legionella sainthelensi ATCC 35248* | 108 | 313 | 171.1 | 3,591 | 33,708 | 0.454 |
| pEK204 | *Prevotella pallens ATCC 700821* | 106 | 435 | 169.9 | 2,860 | 30,096 | 0.609 |

**Table S5**. Key data sample-plasmid pairs with a high excess fraction of PPIs (>0.2) and >100 interacting sample genes and >100 PPIs between the plasmid proteins and the other sample proteins for pEK204 (86 unique proteins), pEK499 (84 unique proteins), pEK516 (55 unique proteins) and pEC958 (44 unique proteins). The table is sorted by the number of interacting proteins.

| **Gene** | **Frequency in 316** | **Frequency in 4,377** | **Name** |
| --- | --- | --- | --- |
| *pilA* | 0.0% | 8.8% | Type IV major pilin protein |
| *pilI* | 19.0% | 1.9% | Purine-binding chemotaxis protein |
| *pilJ* | 21.5% | 6.9% | Transmembrane chemoreceptor (methyl-accepting chemotaxis protein) |
| *pilK* | 3.8% | 0.0% | Chemotactic methyltransferase pilus protein |
| *pilL* | 11.1% | 1.0% | Encode outer membrane lipoprotein |
| *pilM* | 46.2% | 6.3% | Biogenesis of type IV pili |
| *pilN* | 41.8% | 18.4% | Encode outer membrane lipoprotein, secretin |
| *pilO* | 40.8% | 11.8% | Outer membrane, biogenesis of thin pilus |
| *pilP* | 34.2% | 11.5% | Type IV pilus inner membrane component |
| *pilQ* | 62.0% | 9.4% | Biogenesis of type IV pili |
| *pilR* | 22.2% | 0.7% | sigma-54-dependent Fis family transcriptional regulator protein |
| *pilS* | 19.9% | 1.2% | PAS domain-containing sensor histidine kinase |
| *pilT* | 62.0% | 7.7% | Type IV pilus twitching motility protein (retraction of pilus ATPases) |
| *pilU* | 19.3% | 1.7% | Type IV pilus twitching motility protein (retraction of pilus ATPases) |
| *pilV* | 23.4% | 8.9% | Type IV pilus modification protein (recombinase) |
| *pilW* | 0.0% | 1.5% | Type IV pilus biogenesis factor |

**Table S6**. The frequencies of the 14 pEK204-related *pil* genes that were present in 316 bacterial samples on StringDB.

| Name | *fimB* | *fimE* | *fimA* | *fimC* | *fimD* | *fimF* | *fimH* | *fimI* | Total |
| --- | --- | --- | --- | --- | --- | --- | --- | --- | --- |
| *papB* | K12 |  |  |  |  |  |  |  | 1 |
| *papA* |  |  | K12 | C, K12 | C | C, K12 | K12 |  | 5 |
| *papC* |  |  | C, K12 | C, K12 |  | C, K12 | K12 | C | 5 |
| *papD* |  |  | C, K12 | K12 | C | C, K12 | K12 | C | 6 |
| *papF* |  | K12 | K12 | K12 |  | K12 | K12 |  | 5 |
| Total | 1 | 1 | 4 | 4 | 2 | 4 | 4 | 2 | 22 |

**Table S7**. The pairwise PPIs between *fim* and *pap* genes observed for *E. coli* K12 MG1655 (Figure S46) and *E. coli* CFT073 (Figure S47). The presence of a PPIs is marked as “K12” for *E. coli* K12 MG1655, and by “C” for *E. coli* CFT073. 22 PPIs inter-operon PPIs were found: 18 for *E. coli* K12 MG1655 and 11 for *E. coli* CFT073. For *E. coli* K12 MG1655, *papB* stands for *atpD*, *papA* stands for *atpA*, *papC* stands for *atpG*, *papD* stands for *atpB*, and *papF* stands for *atpF*. The PPI pattern for *E. coli* CFT073’s two *pap* operons were identical, and so only one representative is shown here.

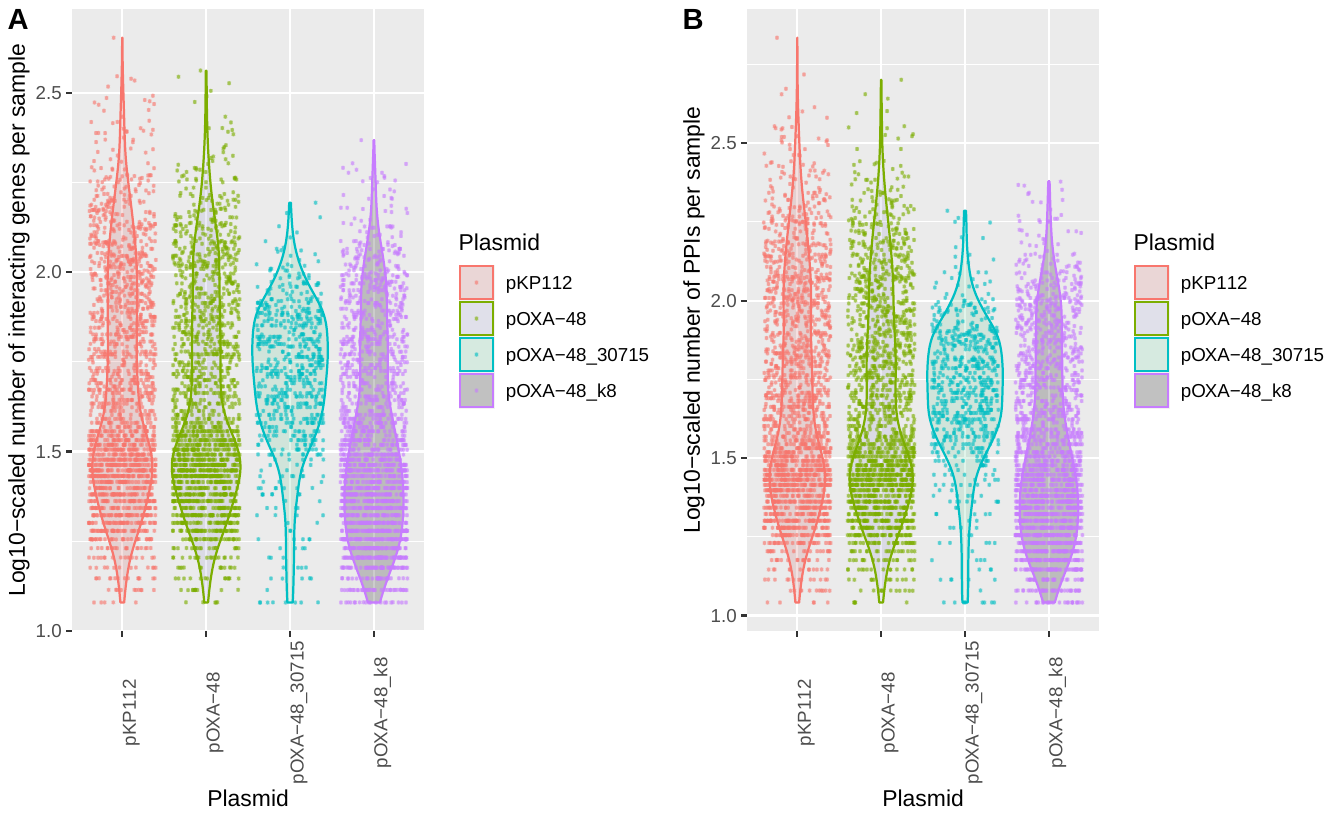

**Figure S1**. The log10-scaled numbers of (A) interacting proteins per sample and (B) PPIs per sample of the pOXA-48 isoforms pKP112 (red-grey), pOXA-48 (green-grey), pOXA-48_30715 (cyan-grey) and pOXA-48_k8 (mauve-grey) using data from 6,181 samples’ plasmids with 10+ interacting proteins and 10+ PPIs each. Plasmid pOXA-48_30715 had relatively higher numbers of interacting sample proteins and PPIs.

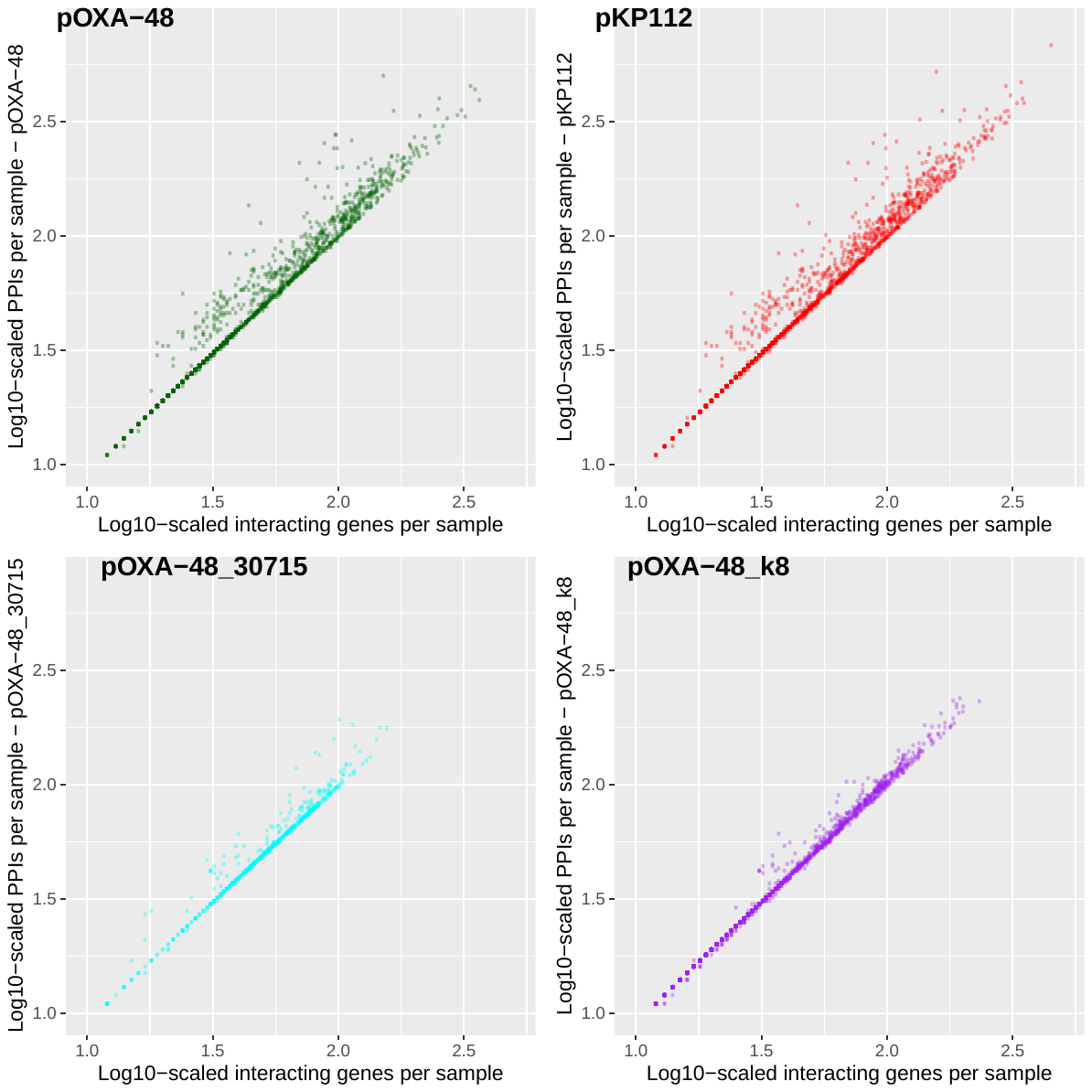

**Figure S2**. The log10-scaled numbers of PPIs per sample (y-axis) relative to the log10-scaled numbers of interacting proteins per sample (x-axis) for each of the four pOXA-48 isoforms (top left) pOXA-48 (green), (top right) pKP112 (red), (bottom left) pOXA-48_30715 (cyan) and (bottom right) pOXA-48_k8 (purple). The unscaled numbers of interacting proteins and PPIs had r=0.963 for pKP112 compared to r=0.957 for pOXA-48, r=0.962 for pOXA-48_30715 and r=0.995 for pOXA-48_k8. The data shown is for 6,181 sample-plasmid pairs with 10+ interacting proteins and 10+ PPIs each.

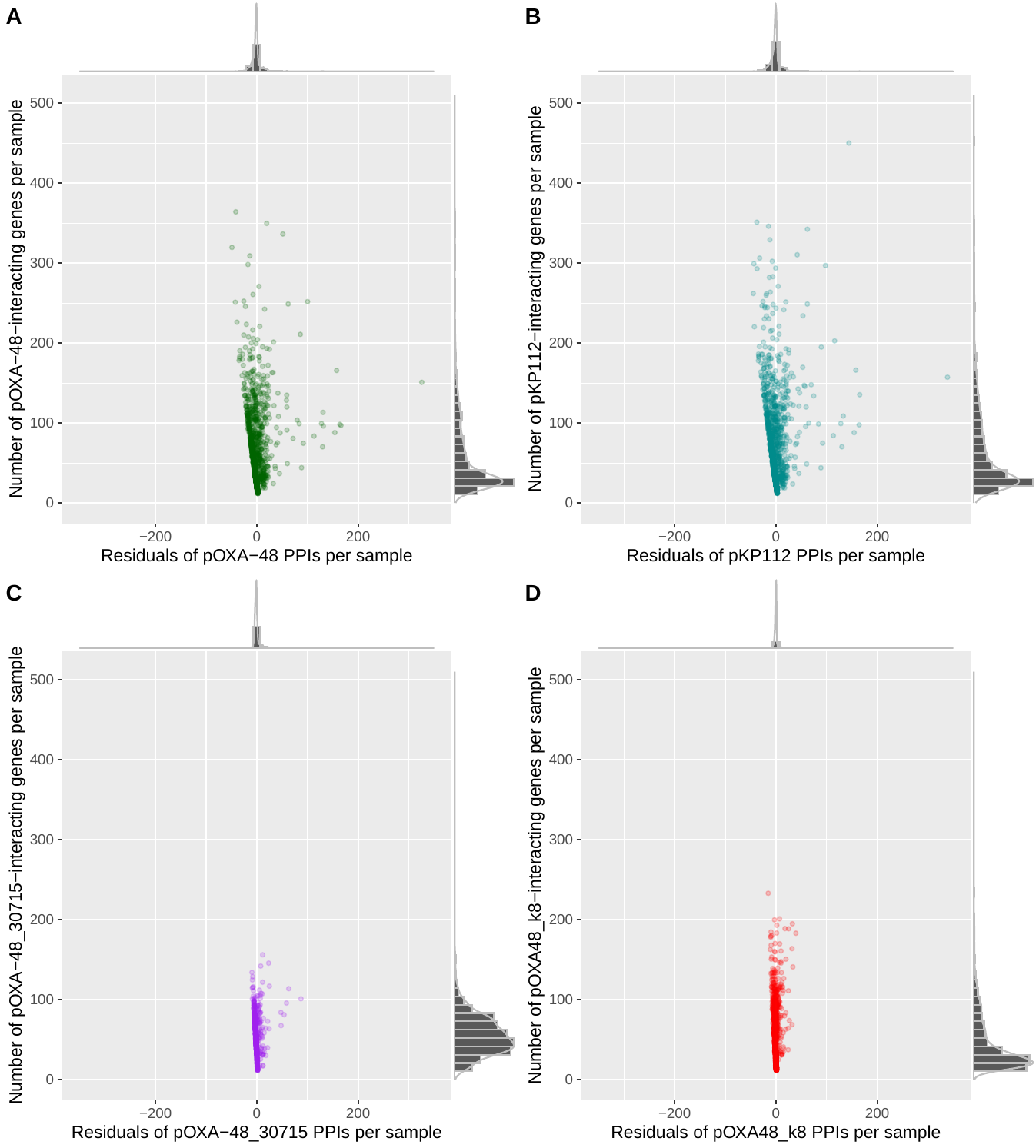

**Figure S3**. The residuals of PPIs with each plasmid per sample (x-axis) versus by the number of sample proteins interacting with each plasmid (y-axis) for the four pOXA-48 isoforms. The plasmids were (A) pOXA-48 (green), (B) pKP112 (cyan), (C) pOXA-48_30715 (purple), and (D) pOXA-48_k8 (red).

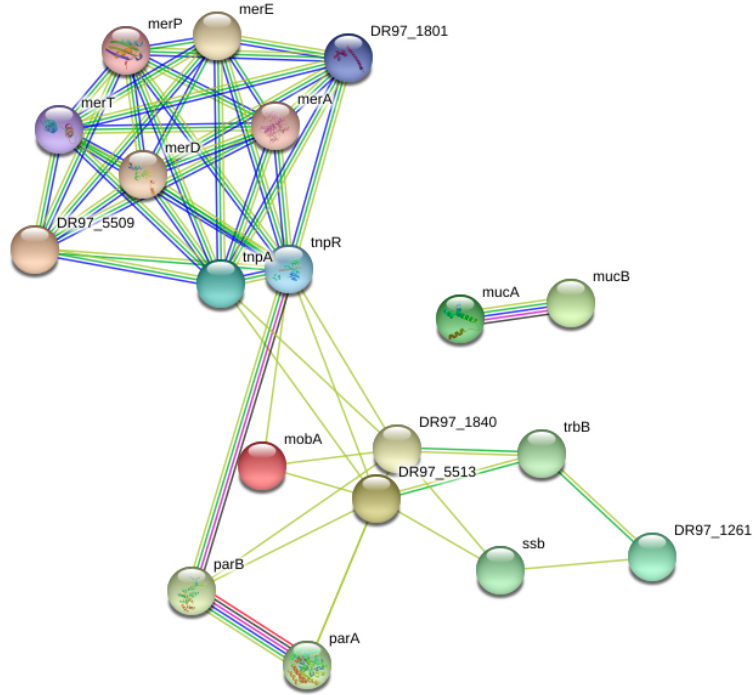
**Figure S4**. A visualisation of a PPI network for *Pseudomonas aeruginosa*’s 19 proteins present on pKP112, which had 55 PPIs. These genes were mainly associated with the *mer* and *muc* operons, the parA-parB partitioning genes, transposon genes, and the *tra* region.

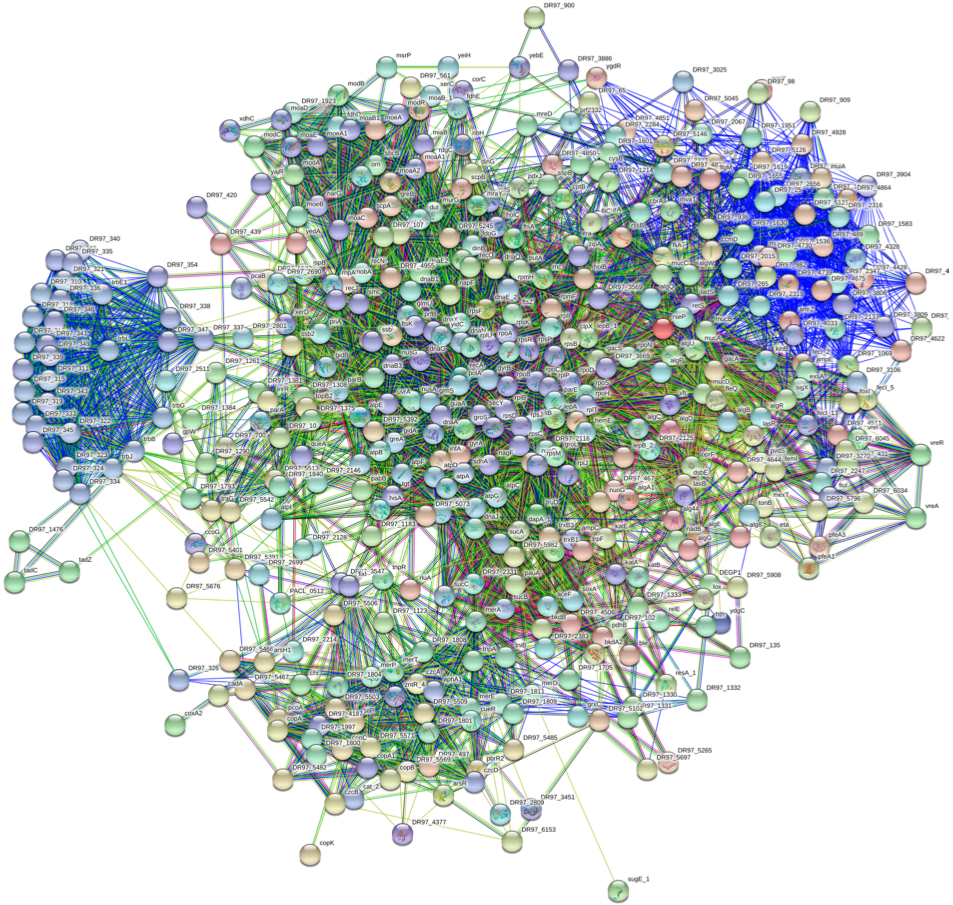

**Figure S5.** A visualisation of a PPI network showing pKP112-related PPIs interacting with 450 of *Pseudomonas aeruginosa*’s proteins.

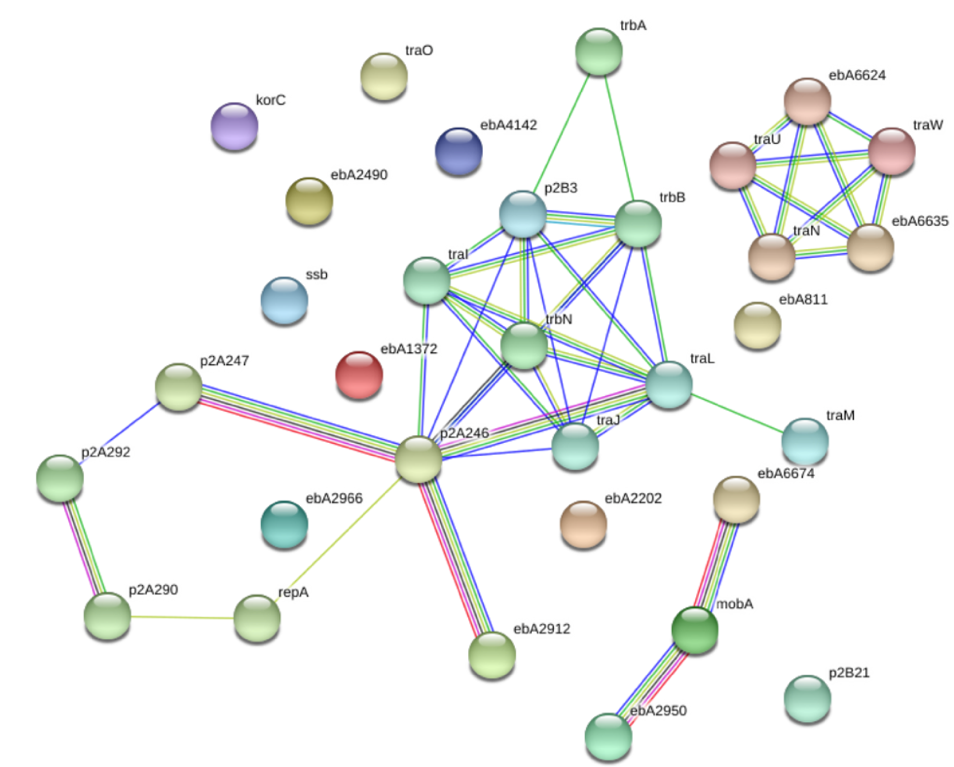
**Figure S6**. A visualisation of the within-plasmid PPI network for *Aromatoleum* *aromaticum* EbN1’s 32 proteins present on pOXA-48, which had 42 PPIs. These genes were mainly associated with the stability genes, *mob* transfer genes, and the *tra* region.

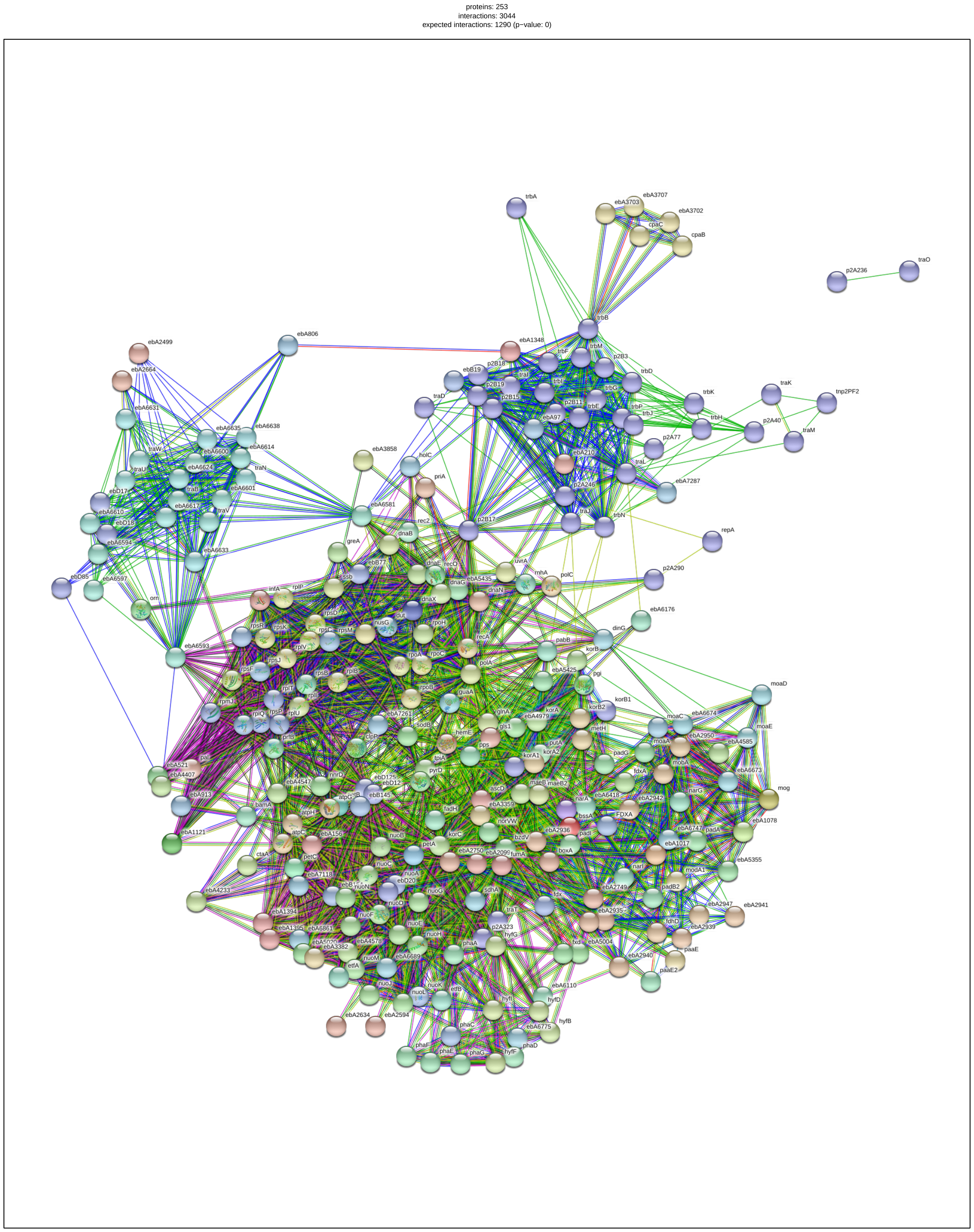

**Figure S7.** A visualisation of a PPI network showing 253 *Aromatoleum* *aromaticum* EbN1 proteins with which pOXA-48 was predicted to interact.

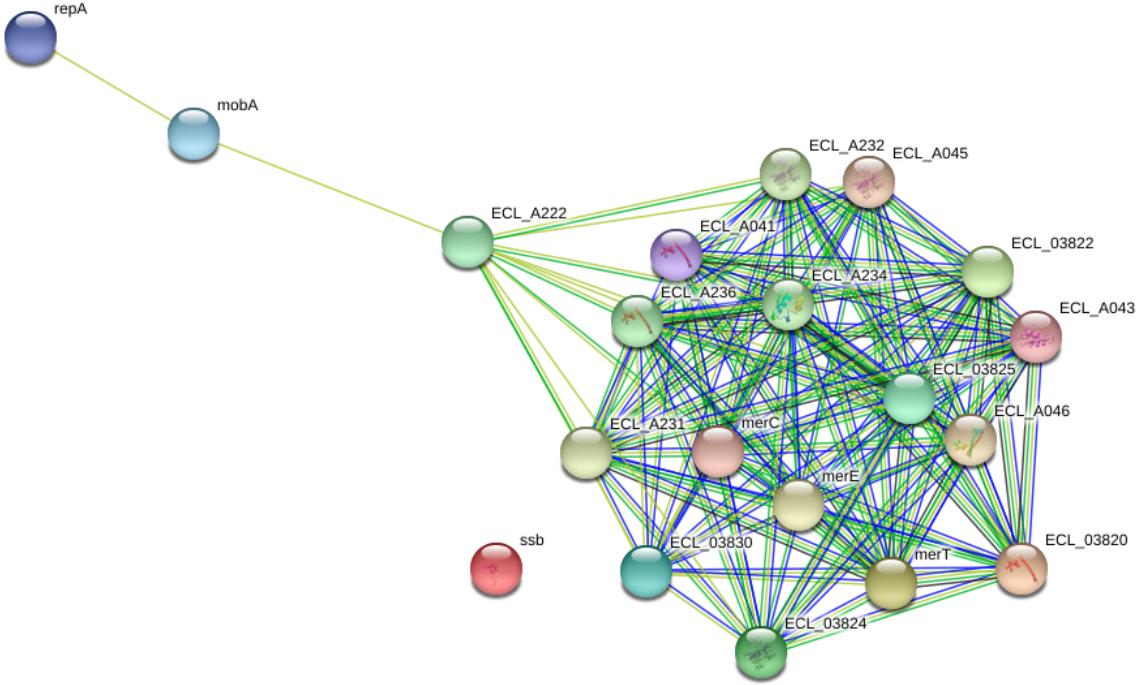

**Figure S8**. A visualisation of a PPI network for *Enterobacter cloacae subsp. cloacae* ATCC13047‘s 12 proteins present on pKP112, which had 121 PPIs. These genes were mainly associated with the *mer* operon.

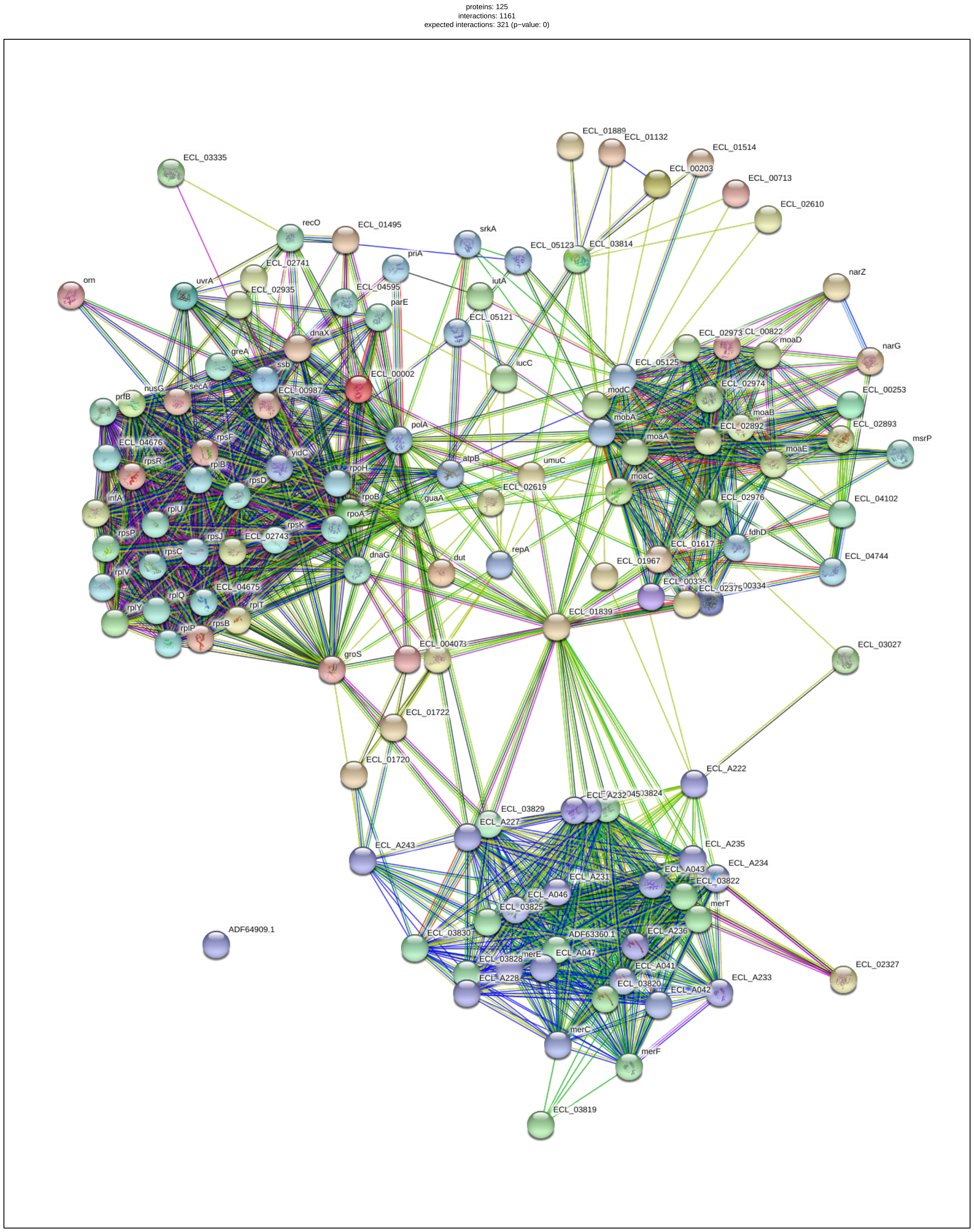

**Figure S9.** A visualisation of a PPI network showing 125 *Enterobacter cloacae subsp. cloacae* ATCC13047 proteins with which pKP112 was predicted to interact.

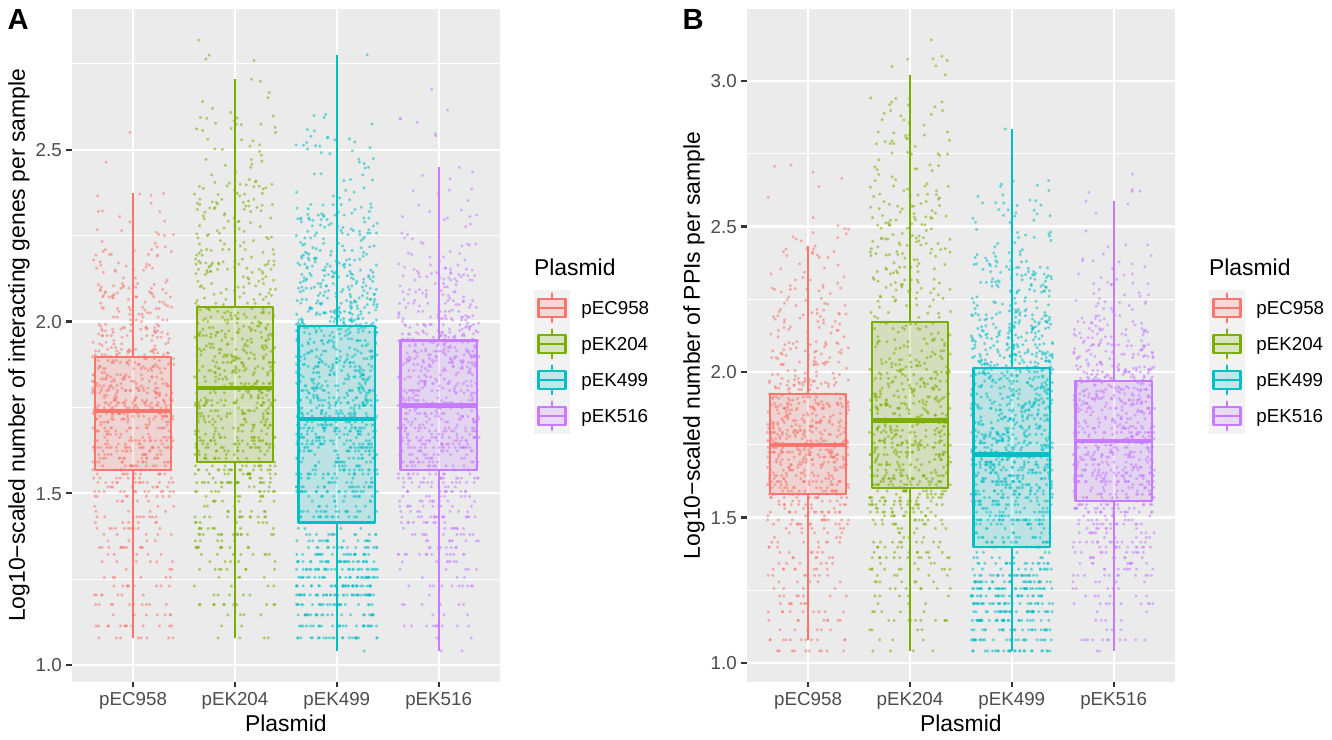

**Figure S10**. The log10-scaled numbers of (A) interacting genes per sample and (B) PPIs per sample of plasmids pEC958 (red), pEK204 (green), pEK499 (cyan) and pEK516 (mauve). The data shown is for 3,828 sample-plasmid combinations with 10+ interacting genes and 10+ PPIs each. Plasmid pEK204 had relatively higher numbers of interacting sample genes and PPIs. Among the 4,419 samples, 968 had interacting genes for pEC958, 901 for pEK204, 892 for pEK499 and 891 for pEK516.

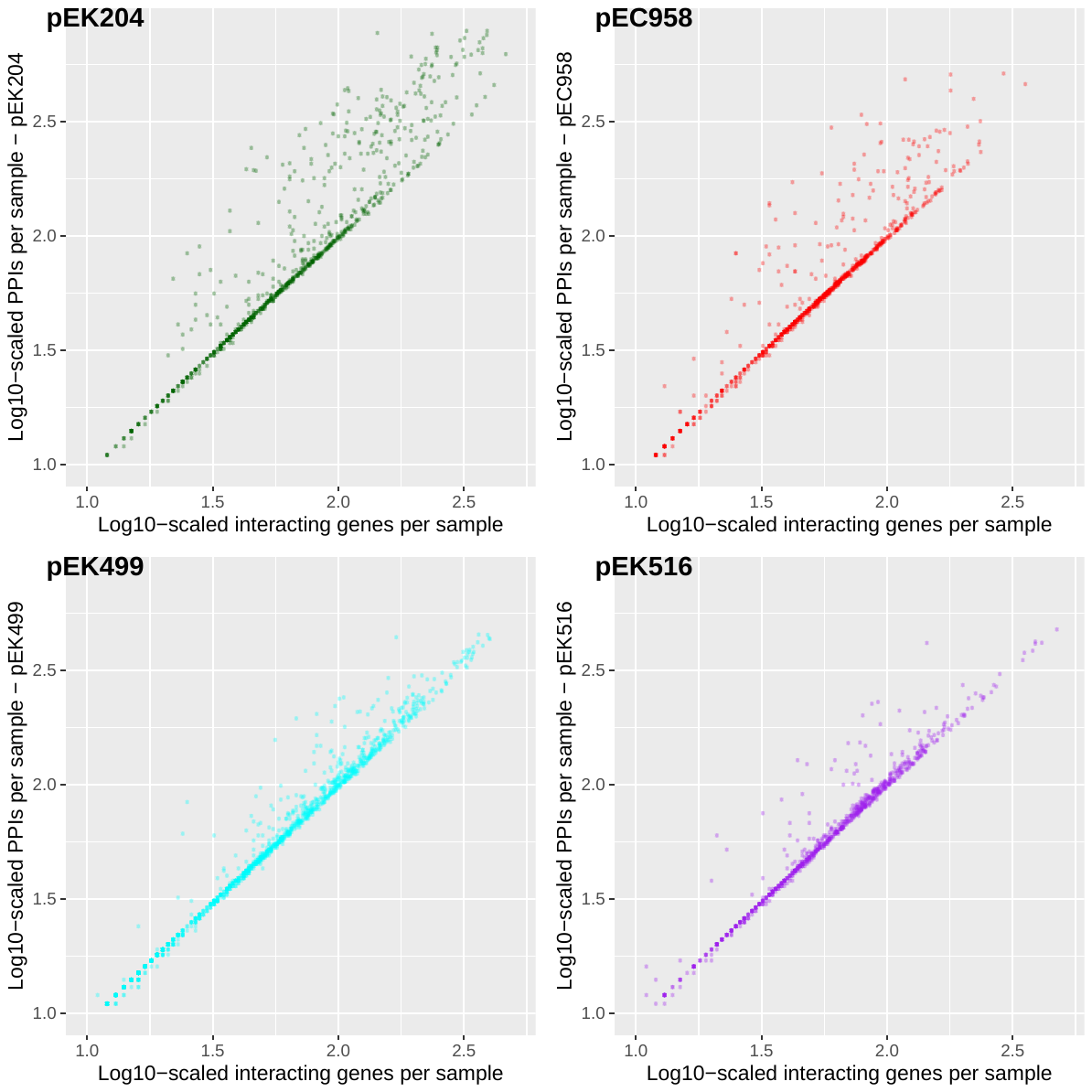

**Figure S11**. The log10-scaled numbers of PPIs per sample (y-axis) relative to the log10-scaled numbers of interacting proteins per sample (x-axis) for each of the four pOXA-48 isoforms (top left) pEK204 (green), (top right) pEC958 (red), (bottom left) pEK499 (cyan) and (bottom right) pEK516 (purple). The unscaled numbers of interacting proteins and PPIs had r=0.895 for pEK204 compared to r=0.853 for pEC958, r=0.980 for pEK499 and r=0.955 for pEK516. The data shown is for 3,828 sample-plasmid pairs with 10+ interacting proteins and 10+ PPIs each.

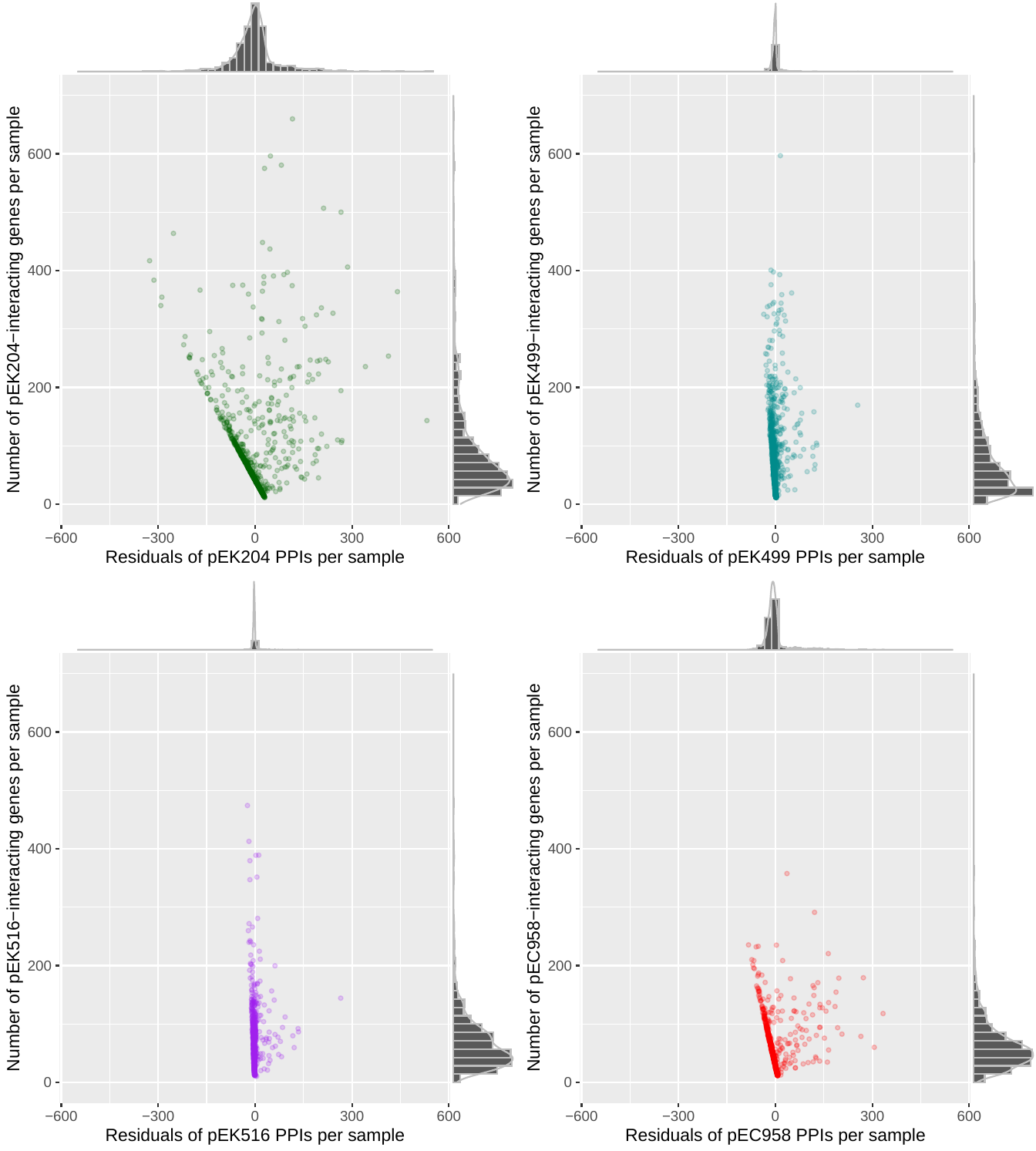

**Figure S12**. The residuals of PPIs with each plasmid per sample (x-axis) versus by the log10-scaled number of sample proteins interacting with each plasmid (y-axis). The plasmids were (A) pEK204 (green), (B) pEK516 (purple), (C) pEK499 (blue), and (D) pEC958 (red).

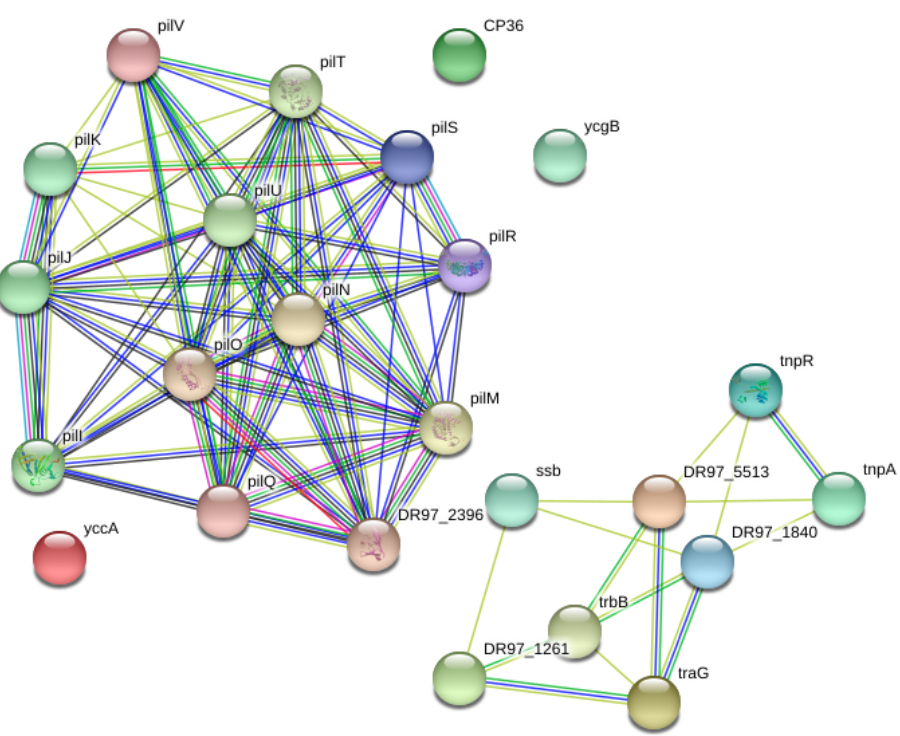

**Figure S13**. A visualisation of a PPI network for *Pseudomonas aeruginosa*’s 24 proteins present on pEK204, which had 84 PPIs. These genes were mainly associated with the *pil* operon (12 genes, including *pilJ-V*) and the *tra* region: *pilI*-*pilV* (excluding *pilP*).

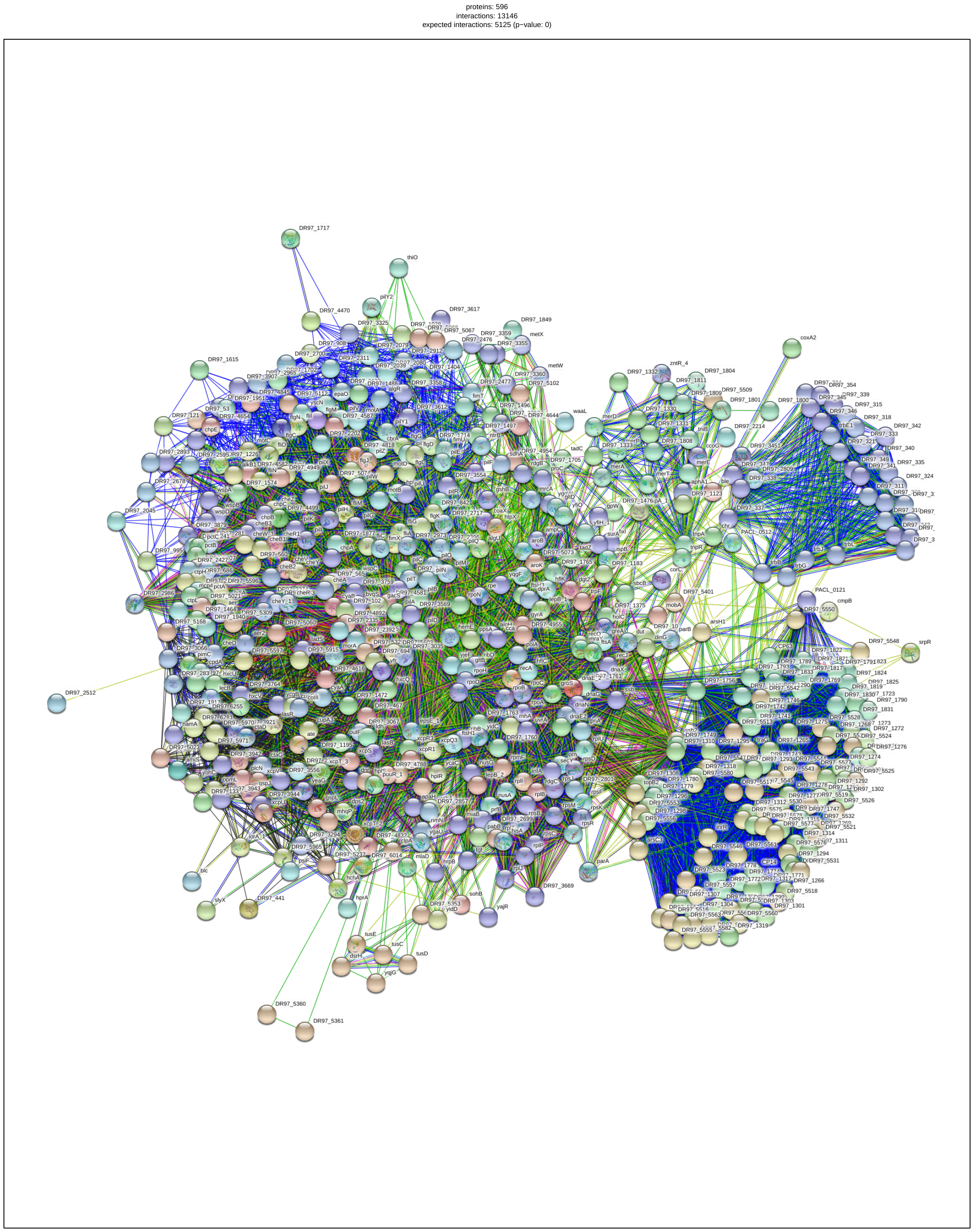

**Figure S14.** A visualisation of a PPI network showing the 596 *Pseudomonas aeruginosa* proteins with which pEK204 was predicted to interact.

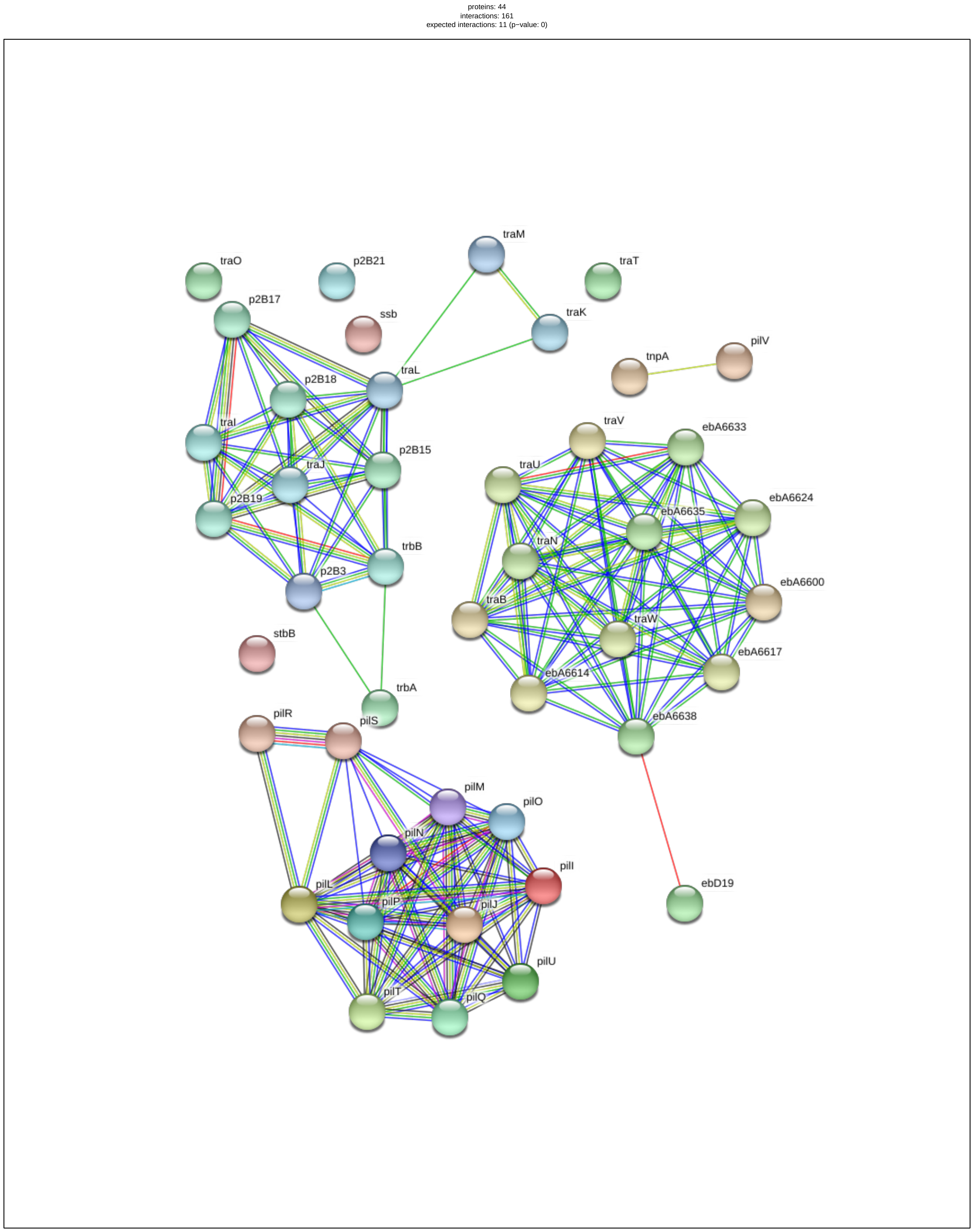
**Figure S15**. A visualisation of a PPI network for *Aromatoleum* *aromaticum* EbN1’s 44 proteins on pEK204, which had 161 PPIs. These genes were mainly associated with the *pil* operon and *tra* region. 12 of these genes were from the *pil* operon: *pilI*-*pilU* (excluding *pilK*).

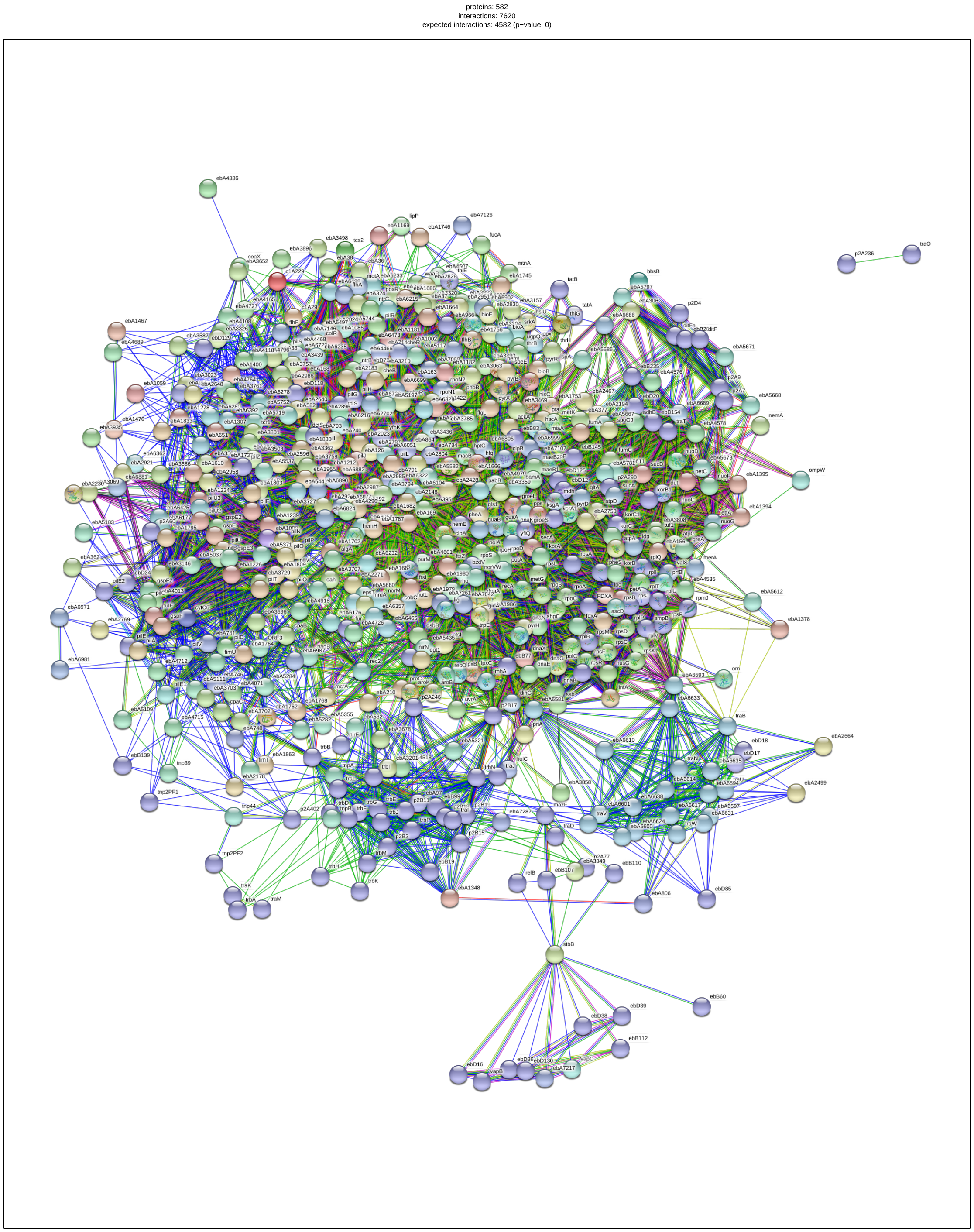

**Figure S16.** A visualisation of a PPI network showing the 582 *Aromatoleum* *aromaticum* EbN1 proteins with which pEK204 was predicted to interact.

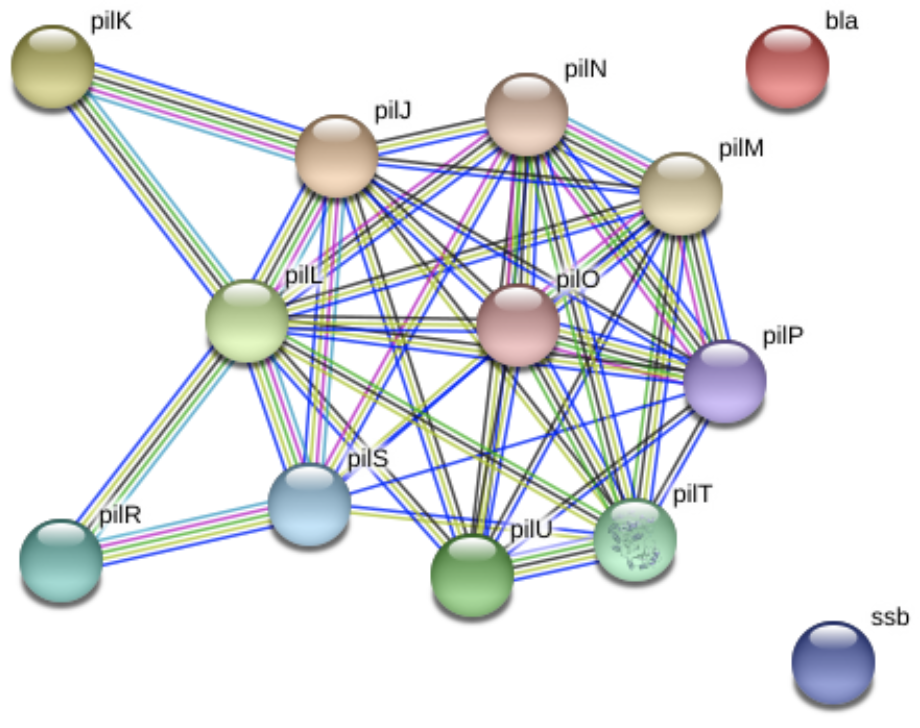

**Figure S17**. A visualisation of a PPI network for *Cellvibrio sp. BR*'s 13 proteins present on pEK204, which had 39 PPIs. These genes were mainly associated with the *pil* operon (11 present): *pilJ*-*pilU* (excluding *pilQ*).

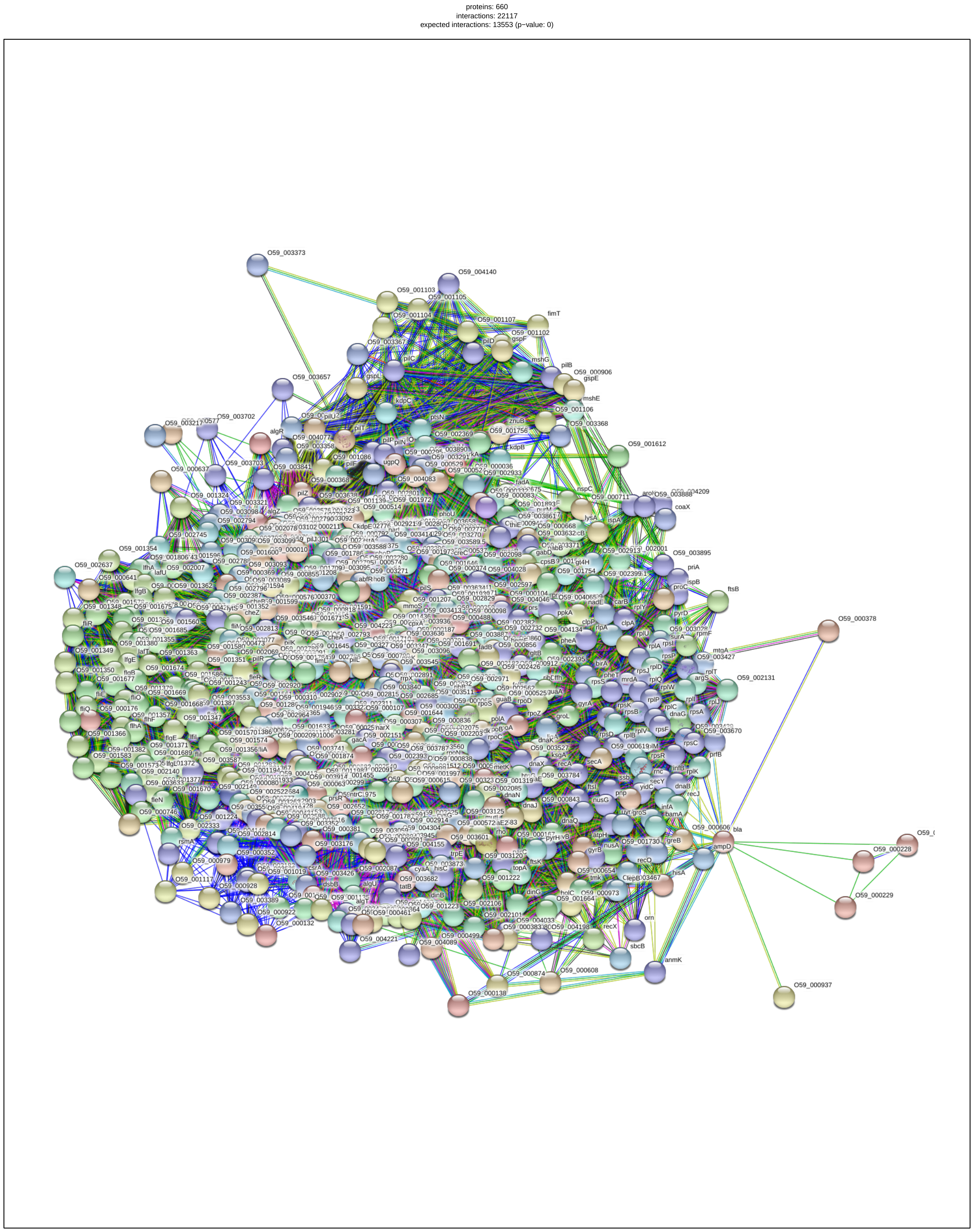

**Figure S18.** A visualisation of a PPI network showing the 660 *Cellvibrio sp. BR* proteins with which pEK204 was predicted to interact.

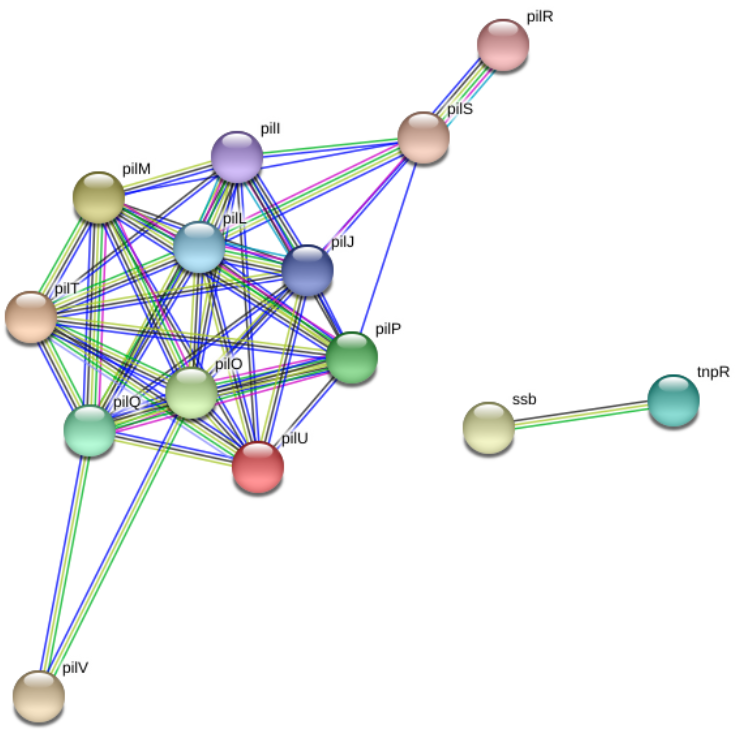

**Figure S19**. A visualisation of a PPI network for *Rubrivivax gelatinosus* IL144’s 14 proteins on pEK204, which had 46 PPIs. 12 of these genes were from the *pil* operon: *pilI*-*pilV* (excluding *pilK* and *pilN*).

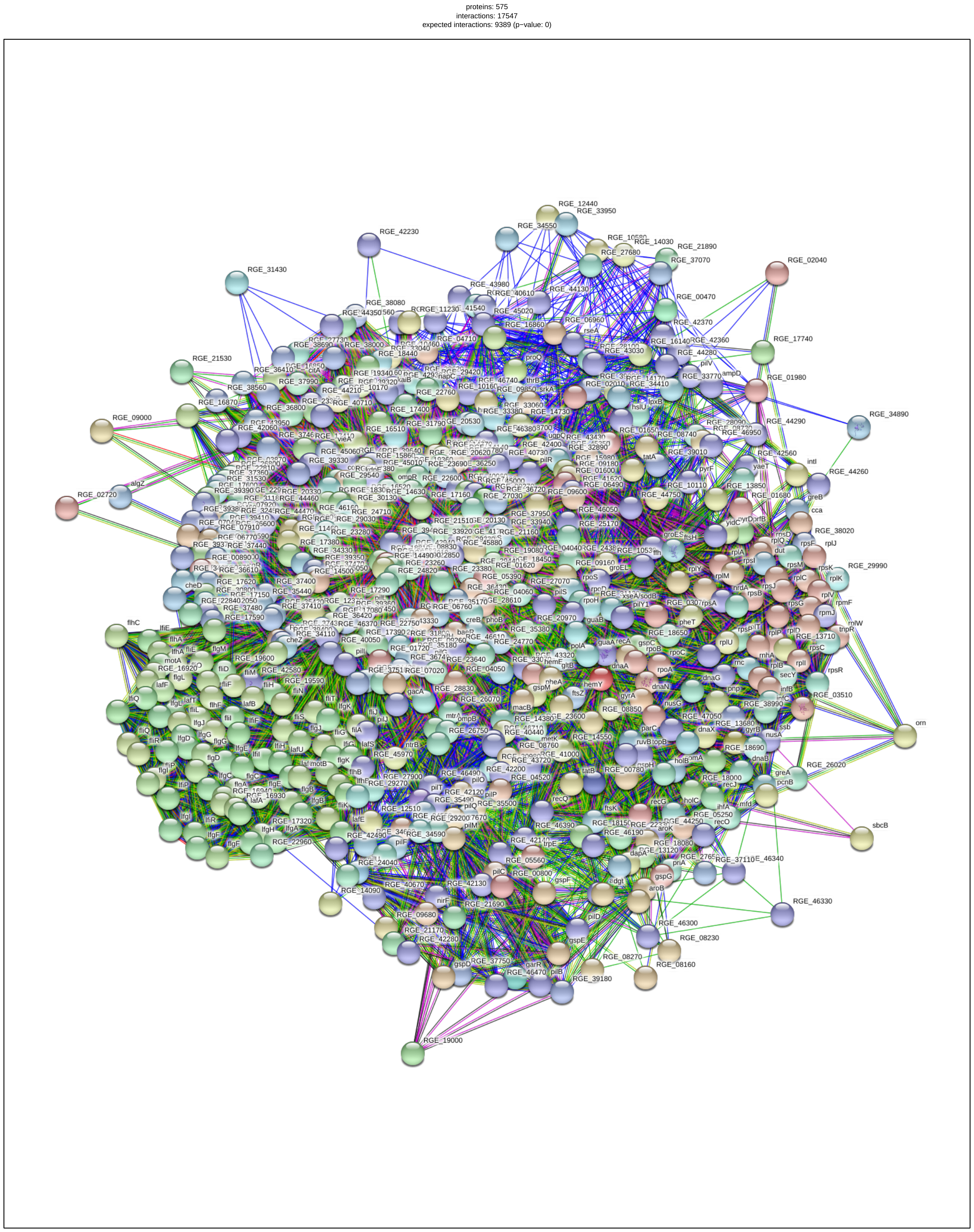

**Figure S20.** A visualisation of a PPI network showing the 507 *Rubrivivax gelatinosus* IL144 proteins with which pEK204 was predicted to interact.

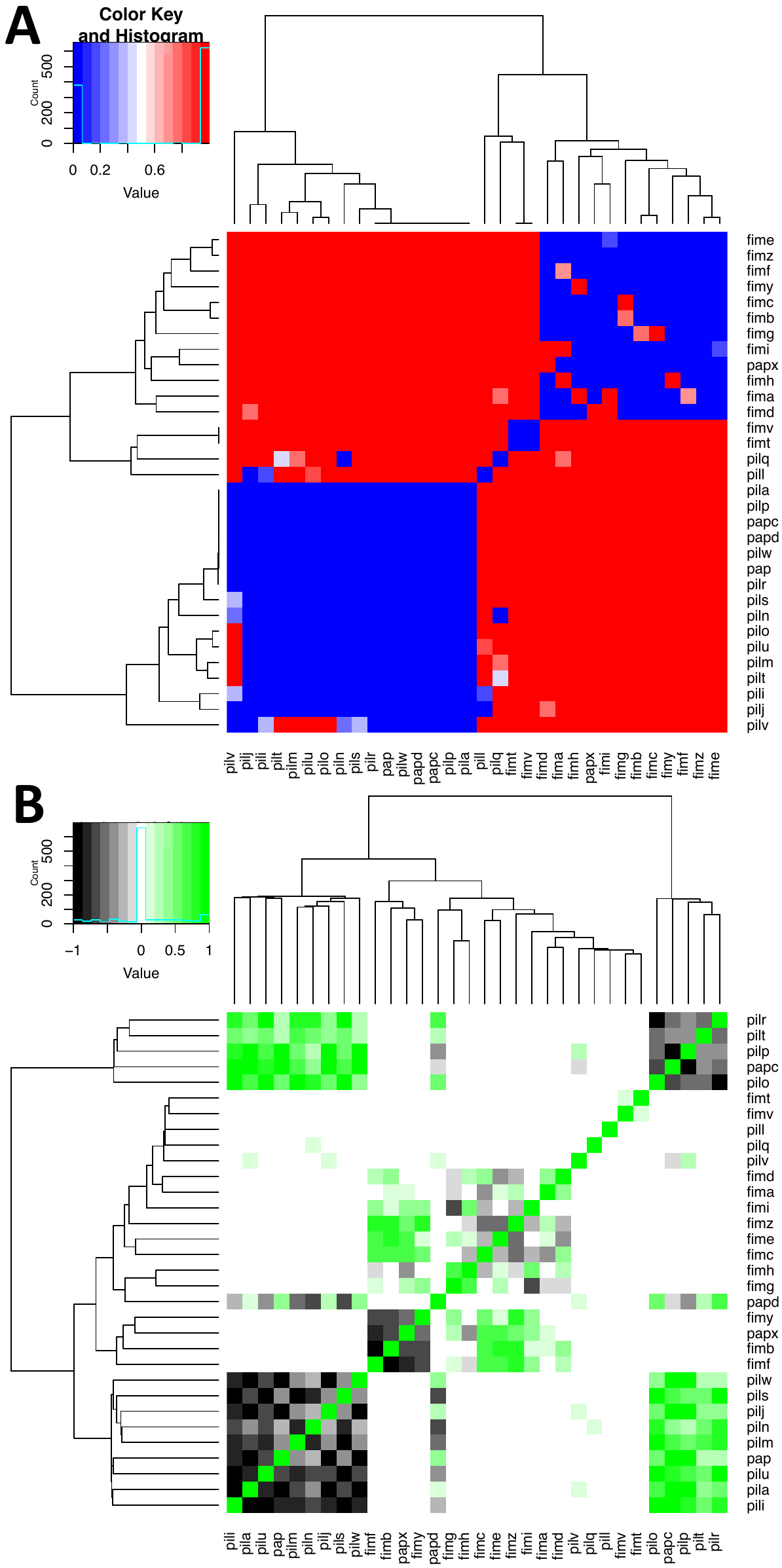
**Figure S21.** The pairwise association based on hierarchical clustering between known *fim, pap* and *pil* genes in bacteria as presence-absence data represented by (A) their correlation coefficient p values and (B) their Pearson correlation coefficients. Both heatmaps are sorted based on similarity on the x- and y-axes. Clustering based on the protein-protein interaction rates per genes showed an identical pattern. In (A) blue indicates small p values, and white to red indicates large ones. All p values were corrected for multiple testing using the Benjamini-Hochberg approach. In (B) black indicates negative correlation coefficients, and green positive ones. This showed three correlated modules, two associated with the *fim* genes and *papX* but excluding *fimT* and *fimV*, and the other two associated with the *pil* and *pap* genes but excluding *pilQ*, *pill* and *pilV*. Thus, *fimT*, *fimV*, *pilQ*, *pilL* and *pilV* made up one largely uncorrelated module. Within the *fim*-related module the genes were positively correlated with one another, except for *fimB*, *fimF,* *fimY* and *papX* (aka transcriptional repressor *mprA*), which were negatived correlated with one another, but were positively associated with *fimA*, *fimC*, *fimD*, *fimE*, *fimZ* and *fimI*. Within the *pil*/*pap*-related module, the genes were positively correlated with one another, although there were two blocks of negatively correlated genes: one for *pilO*, *pilP*, *pilR*, *pilT* and *papC*, and the other for *pap*, *papD*, *pilA*, *pili*, *pilJ*, *pilM*, *pilN*, *pilS*, *pilU*, *pilW* and *pilV*.

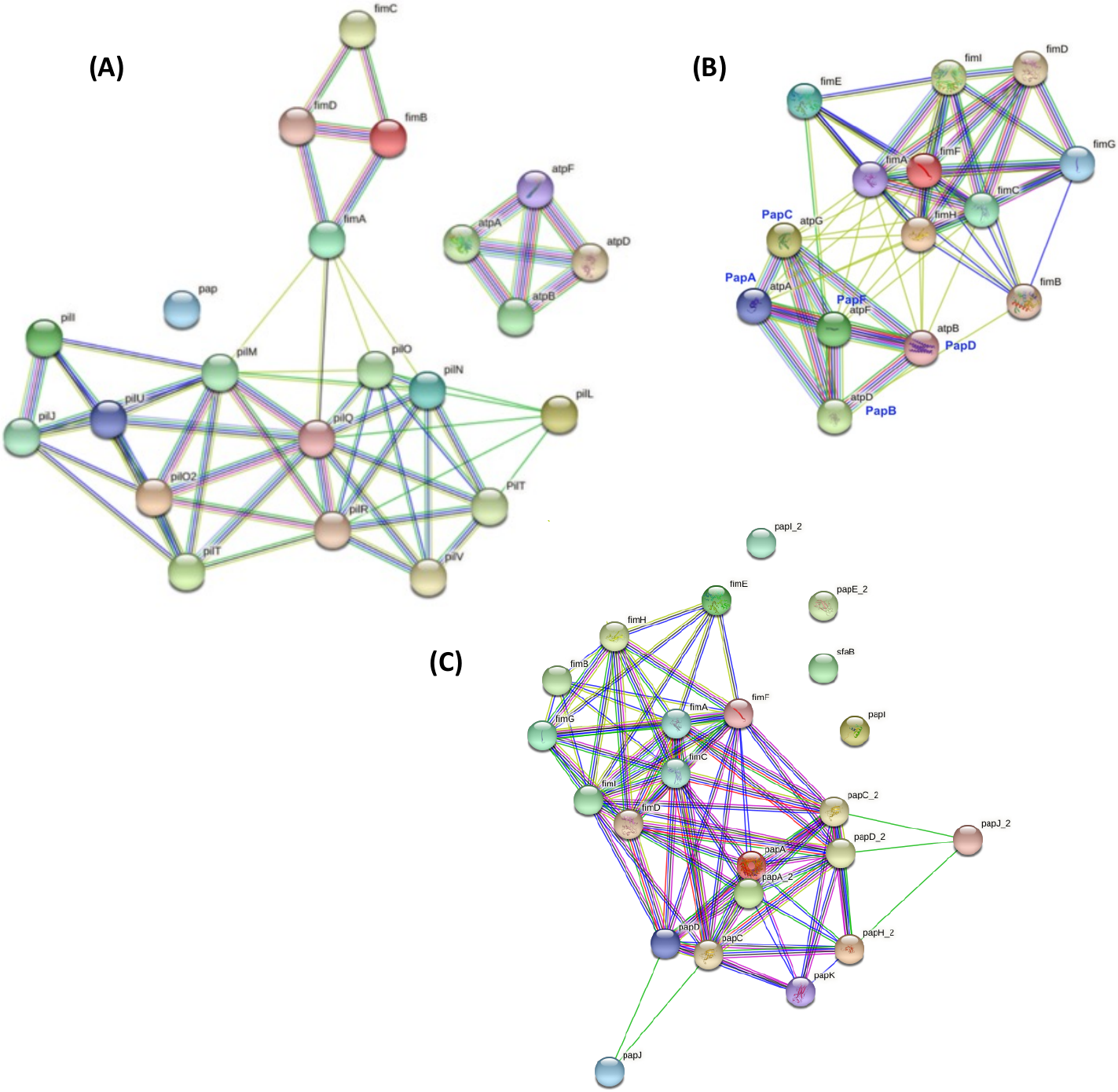

**Figure S22**. Visualisation of PPI networks for (A) *Ralstonia eutropha* H16, (B) *E. coli* K12 MG1655 and (C) *E. coli* CFT073. (A) *Ralstonia eutropha* H16 had 22 proteins matched *fim*, *pap* or *pil* operon genes: 13 of these genes were from the *pil* operon and four from the *fim* operon. These had 68 PPIs, including for FimA with PilM, PilN, PilO and pilQ. (B) *E. coli* K12 MG1655 had 14 proteins whose names matched *fim*, *pap* or *pil* operon genes: five of these genes were from the *pap* operon and nine from the *fim* operon. These had 60 PPIs. (C) *E. coli* CFT073’s 23 proteins whose names matched *fim*, *pap* or *pil* operon genes: five of these genes were from *pap* operon 1, five were from *pap* operon 2 and 11 were from the *fim* operon. These had 84 PPIs. Genes matching the *apt* operon were not examined. The gene name synonyms included: F7-2 as PapA and FooG as PapG in *E. coli* CFT073; and FooB as PapB in 536; and the Pap proteins for *E. coli* 536 were: ECP_4533 (PapG), ECP_4536 (PapK), ECP_4538 (PapD), ECP_4539 (PapC), ECP_4540 (PapH), ECP_4541 (PapA) and ECP_4532 (PapE/PapF); and for *E. coli* K12 MG1655, atpG was PapC, atpA was PapA, aptH was PapE, aptC was PapG, aptE was PapH, atpF was PapF, atpB was PapD and atpD was PapB. Other synonyms were the normal name followed by an underscore and/or a digit.
